# The bipolar-to-multipolar transition of dendrite extension of cerebellar granule neurons requires MTCL2 to link the Golgi apparatus to the microtubule cage

**DOI:** 10.1101/2024.09.12.612747

**Authors:** Mari Minekawa, Atsushi Suzuki

## Abstract

The dynamic regulation of neuronal polarity is essential for the formation of neural networks during brain development. Primary cultures of rodent neurons recapitulate several aspects of this polarity regulation, providing valuable insights into the molecular mechanisms underlying axon specification, dendrite formation, and neuronal migration. However, the process by which the preexisting bipolarity of migrating neurons is disrupted to form multipolar dendrites remains to be elucidated. In this study, we demonstrate that MTCL2, a microtubule-crosslinking protein associated with the Golgi apparatus, plays a crucial role in this type of polarity transformation observed during the differentiation of cerebellar granule neurons (CGNs). MTCL2 is highly expressed in CGNs and gradually accumulates in dendrites as the cells develop polarity. MTCL2 knockdown resulted in the generation of longer and fewer dendrites by suppressing the bipolar-to-multipolar transition of dendrite extension observed in the normal polarization process. During this process, the Golgi apparatus shifts from the base of the preexisting bipolar neurites to the lateral or apical side of the nucleus in the cell body. There, it forms a close association with the microtubule cage that wraps around the nucleus. The resulting upward extension of the Golgi apparatus is tightly coupled with the randomization of its position in the x-y plane. Knockdown and rescue experiments demonstrated that MTCL2 promotes these Golgi position changes in a microtubule- and Golgi-binding activity-dependent manner. These results suggest that MTCL2 promotes the development of multipolar short dendrites by sequestering the Golgi apparatus into the microtubule cage from the base of the preexisting neurite.

**Significance Statement:** Previous studies have shown that neurons polarize the localization of the Golgi apparatus, thereby facilitating the development of asymmetrically elongated dendrites. In contrast, the present study demonstrated that to symmetrically extend multiple dendrites, cerebellar granule neurons sequester the Golgi apparatus within a microtubule cage surrounding the nucleus. This represents a novel mechanism through which neurons suppress polarized vesicle transport. These findings also reveal an entirely novel function of microtubule cages, which have only been reported for their role in neuronal cell migration. The present study also revealed the involvement of the novel microtubule-regulating protein MTCL2, which links the Golgi apparatus to microtubules, and the LINC complex, which links microtubules to the nuclear membrane, in this process.

## Introduction

Neurons are highly polarized cells in the human body that develop two types of functionally distinct processes: a long, thin axon and short dendrites. To establish elaborate neuronal circuits in the brain, developing neurons gradually establish neuronal polarity while migrating to appropriate regions.

Neuronal polarization is intrinsically programmed and can be recapitulated *in vitro* to some extent, even without specific cell-to-cell contacts or extracellular matrix molecules (Tahirovic and Bradke, 2009). Rodent hippocampal neurons are the most intensively studied model system for investigating neuronal polarity (Dotti et al., 1988; Fletcher and Banker, 1989). Cells isolated from late embryos first develop a multipolar morphology with several short neurites that exhibit rapid elongation and retraction. Remarkably, one of these initial neurites begins to grow rapidly and becomes a thin, long axon 1-2 days after plating. The remaining short neurites then develop into dendrites. Most studies have focused on the initial event through which neurons break the initial multipolarity, as similar axon specification processes have been observed *in vivo* in the developing neocortex (Miyata et al., 2004; Noctor et al., 2004; Takano et al., 2015).

Dissociation cultures of cerebellar granule neurons (CGNs) reflect the characteristic developmental process of this type of neurons *in vivo,* and provide an additional model for studying neuronal development (Tahirovic and Bradke, 2009; Powell et al., 1997; Zmuda and Rivas, 1998; Solecki et al., 2006). In contrast to hippocampal neurons, the initial development of CGNs isolated from early postnatal rodents is characterized by the formation of bipolar axons without the elongation of multiple neurites. Subsequently, the cells transform the bipolar axon into a single T-shaped axon by either migrating perpendicular to the initial bipolar axis or by retracting one of the two axons. Finally, they extend short dendrites in multiple directions to complete their final morphology (Powell et al., 1997; Solecki et al., 2006; Tahirovic and Bradke, 2009; Zmuda and Rivas, 1998). Researchers initially employed this distinctive property of CGNs in culture to demonstrate the close correlation between the position of the centrosome and the Golgi apparatus with the site of axon elongation (Zmuda and Rivas, 1998). This culture system was then used to investigate the molecular mechanisms underlying axonogenesis (Arakawa et al., 2003; Oishi et al., 2021) and dendritogenesis (Huang et al., 2007; Litterman et al., 2011; Tan et al., 2013). However, no studies have focused on the unique polarity change, the transition from bipolar to multipolar, exhibited by CGNs, but not by hippocampal neurons. Consequently, the process by which CGNs disrupt preexisting bipolarity to extend multipolar dendrites remains unclear.

MTCL2, formerly known as SOGA1, is a recently identified microtubule-associated protein (Matsuoka et al., 2022). It dimerizes through its central coiled-coil region and interacts with microtubules (MTs) via its C-terminal region. The N-terminal portion of the coiled-coil region also exhibits additional activity to associate with the Golgi membrane through interactions with CLASPs and Giantin. These abilities enable MTCL2 to accumulate MTs around the Golgi apparatus, thereby promoting the lateral assembly of Golgi stacks and forming a compact Golgi ribbon. Using a wound-healing assay with a cultured cell line, Matsuoka et al. (2022) further suggested that these MTCL2 functions are essential for maintaining directional migration.

In this study, we demonstrated that MTCL2 is highly expressed in CGNs *in vivo*. In cultured CGNs, MTCL2 depletion resulted in abnormally polarized cells with longer and fewer dendrites. Several experiments, including live-cell imaging, revealed that MTCL2 is necessary for the bipolar-to-multipolar transition of dendrite extensions, a process observed during the normal development of CGNs. Immunofluorescence analysis and knockdown/rescue experiments have revealed that MTCL2 sequesters the Golgi apparatus from the base of the bipolar neurites to the MT cage that wraps around the nucleus. The importance of the MT cage was also demonstrated by inhibiting the LINC complex, which links the nuclear membrane to MTs. Taken together, these results demonstrate MTCL2’s novel role in promoting the bipolar-to-multipolar transition of dendrite extensions by linking the Golgi apparatus and MTs.

## Results

### MTCL2 shows preferential enrichment in the dendrites of cerebellar granule neurons

A previous study by Matsuoka et al. (2022) showed preferential expression of MTCL2 in the lung, testis, ovary, cerebrum, and cerebellum using western blotting of mouse tissue extracts. High expression of MTCL2 in the cerebellum is also evident in public mRNA expression databases for human and mouse organs (https://www.ebi.ac.uk/gxa/home and https://www.proteinatlas.org), as well as *in situ* hybridization data for the mouse brain (https://mouse.brain-map.org/) (Fig. S1A). We therefore performed an immunostaining analysis of MTCL2 in the mouse cerebellum and found that MTCL2 is preferentially expressed in granule cells and, to a lesser extent, in Bergmann glial cells, but not in Purkinje cells (Fig. 1A; A. Suzuki, unpublished observation). The absence of MTCL2 in Purkinje cells is consistent with the public *in situ* hybridization data for the mouse brain (Fig. S1), whereas it is in sharp contrast to its paralogue, MTCL1, which is expressed in Purkinje cells and plays critical functions (Satake et al., 2017). Based on these results, in subsequent studies, we analyzed MTCL2 functions in cerebellar granule neurons (CGNs) using dissociated cultures of mouse CGNs (Krämer and Minichiello, 2010; Powell et al., 1997).

**Figure 1.**
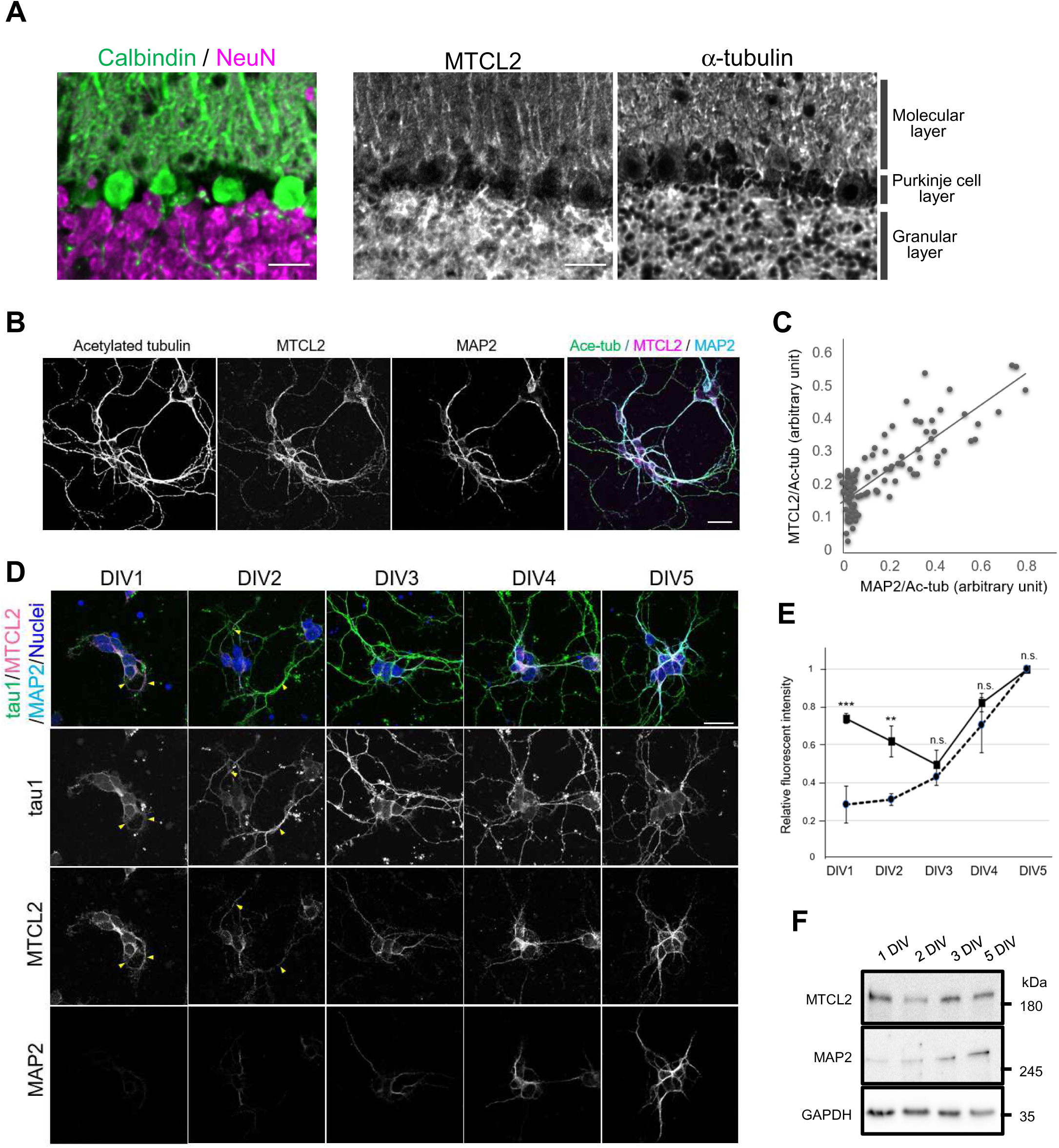
MTCL2 is preferentially enriched in the dendrites of cerebellar granule neurons. (A) Cerebellar sections from 4-week-old mice are stained for calbindin and NeuN (left panel) or MTCL2 and α-tubulin (middle and right panels). Scale bars: 250 μm. (B) Representative images of isolated CGNs cultured for 5 days and stained for acetylated tubulin, MTCL2, and MAP2. Scale bar: 20 μm. (C) Correlation between the immunofluorescence intensities of MTCL2 and MAP2 in acetylated tubulin-positive neurites. A total of 100 neurites from 3 independent experiments were analyzed. (D) Representative images of CGNs at DIV1–DIV5 stained for tau1, MTCL2, and MAP2. In the merged view, nucleus staining is included to clarify the positions of cell bodies. The arrowheads indicate tau1-positive neurites exhibiting MTCL2 signals in immature CGNs. Scale bar: 20 μm. (E) Time-dependent increase in the relative intensities of the MTCL2 (solid line) and MAP2 (dotted line) signals in dendrites. Neurites exhibiting MAP2 signal intensities three times greater than the background were analyzed as dendrites. Data represent the means ± S.D. from 3 independent experiments (totally, more than 67 neurites were analyzed for each DIV stage). The paired t-test was used to examine statistical significance (**p < 0.01; ***p < 0.005; n.s., non-significant). (F) Western blot analysis of MTCL2 and MAP2 expression in primary cultured CGNs from various DIVs. A representative result of 2 independent experiments is presented.

Immunofluorescence analysis of CGNs isolated and cultured for 5 days *in vitro* (DIV5) revealed strong anti-MTCL2 signals in the cell bodies and in some restricted neurites (Fig. 1B). The disappearance of these signals following MTCL2 knockdown (Fig. S2) indicates that they represent the true localization of endogenous MTCL2. Neurites with a strong MTCL2 signal showed a high degree of correlation with the MAP2-positive dendrites (Fig. 1B). In fact, the intensities of MTCL2 and MAP2 staining on the neurites were highly correlated (Fig. 1C; Pearson’s r = 0.83). However, weak MTCL2 signals were also detected in MAP2-negative thin axons (Fig. 1B, C). These results suggest that, while MTCL2 is not restricted to MAP2-positive dendrites, it is preferentially enriched in these dendrites.

Next, we examined the timing of MTCL2 enrichment in dendrites during CGN polarization. To this end, we compared the intensities and patterns of the immunostaining for dephosphorylated tau (tau1), MAP2, and MTCL2 in CGNs cultured for 1–5 days (Mandell and Banker, 1996). Consistent with previous reports, CGNs extended tau1-positive short processes 1 day after seeding (Fig. 1D, arrowheads) (Zmuda and Rivas, 1998). Most of these processes acutely elongated after DIV2 and gradually became axon-like neurites with longer and thinner morphologies. Conversely, MAP2 signals were very weak until DIV3, when they began to be detected in the short neurites. In contrast, MTCL2 signals were strongly detected even at DIV1, at levels comparable to those observed at DIV5 (∼80%). This level of expression was maintained thereafter, with slight decreases at DIV2 and 3 (Fig. 1D,E). High expression of MTCL2 in DIV1 CGNs was also confirmed by western blotting (Fig. 1F). MTCL2 signals in CGNs at DIV1 and 2 were detected in cell bodies and in some tau1-positive initial neurites (Fig. 1D, arrowheads). Around DIV3-4, however, when MAP2-positive neurites appeared, MTCL2 gradually exhibited selective enrichment in these neurites (Fig. 1D). These results suggest that MTCL2 enrichment in dendrites occurs during the late stage of CGN polarization in parallel with MAP2 expression.

### Depletion of MTCL2 resulted in the development of fewer and longer dendrites

These results suggest a possibility that MTCL2 plays an important role in the dendritic development of CGNs. To investigate this possibility, we examined the effects of MTCL2 knockdown in primary cultured CGNs. Eight hours after seeding, we transfected a plasmid expressing shRNA and EGFP into the cells (Fig. S2). As shown in Fig. 2A, CGNs lacking MTCL2 appeared to differentiate normally into neurons with long, thin axons at DIV5. Quantification of axon length and branching revealed no significant differences between MTCL2-depleted cells and control cells (Fig. 2B; Fig. S3A). Furthermore, MTCL2 knockdown did not prevent the appearance of MAP2-positive dendrites in DIV6 CGNs (Fig. 2C). However, detailed analyses revealed that MTCL2 knockdown resulted in fewer dendrites than control cells in terms of both the number of dendrite tips (Fig. 2D) and bases (Fig. S3B). Additionally, the average dendrite length of MTCL2-depleted CGNs (25.0 ± 11.5 µm) was greater than that of control cells (15.4 ± 4.0 µm) (Fig. 2E). Analysis of the length distribution of the total dendrites revealed that the increase in average length was primarily due to the existence of long dendrites in the MTCL2-depleted cells, which were rarely observed in the control cells (Fig. S3C). For instance, the maximum dendritic length of individual CGNs was 35.7 ± 11.5 µm on average for control cells and 54.0 ± 21.6 µm for the MTCL2-depleted cells (Fig. S3D). The proportion of dendrites longer than 40 μm in the total number of dendrites was 3.14 ± 0.13% in control cells and 14.36 ± 0.10% in MTCL2-depleted cells (Fig. 2F). These changes in dendrite numbers and length distribution in MTCL2-depleted cells were reversed by expressing RNAi-resistant mouse MTCL2 (Fig. 2D-F; Fig. S3B-D). These results suggest that MTCL2 depletion leads to abnormal CGN development, resulting in fewer longer dendrites.

**Figure 2.**
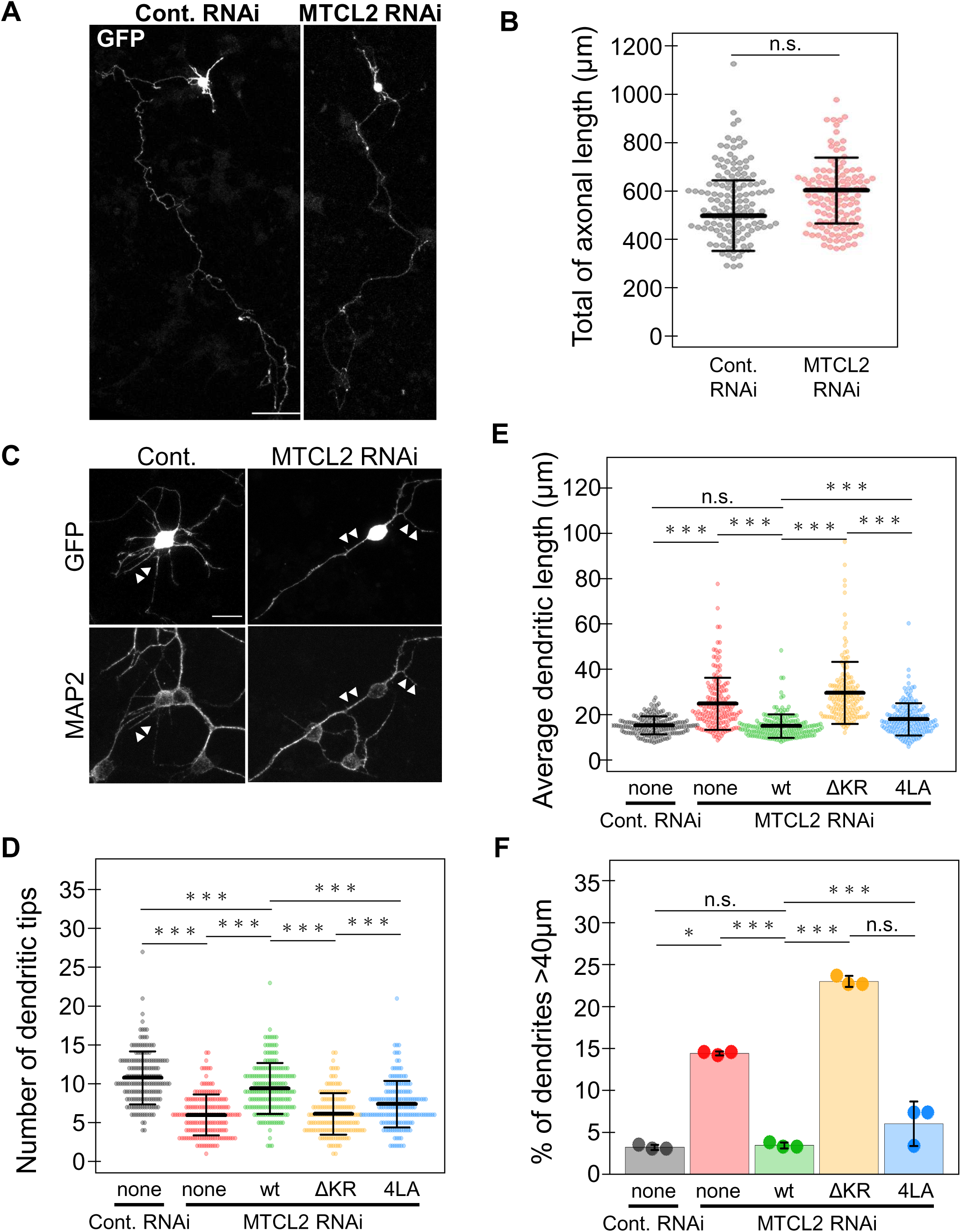
Depletion of MTCL2 resulted in the development of fewer and longer dendrites. (A) Full views of single CGNs at DIV6 visualized by the GFP signal. Scale bar: 50 μm. (B) Distribution of total axonal length of individual CGNs subjected to the indicated RNAi and fixed at DIV6. (C) Dendritic development of DIV6 CGNs transfected with control or MTCL2 knockdown vectors and visualized by GFP and MAP2 signals. The arrowheads indicate the dendrites of GFP-positive cells. Scale bar: 10 μm. (D) Distribution of total number of dendritic tips of individual CGNs subjected to the indicated RNAi/rescue conditions and fixed at DIV6. (E) Distribution of averaged dendritic lengths of the individual CGNs analyzed in (D). (F) Percentage of dendrites longer than 40 μm in all dendrites of the same CGNs analyzed in (D) and (E). Each data point in (B), (D), and (E) represents the result of one neuron in 3 independent experiments. Mean and standard deviations are shown for each scatter plot. The bars in (F) represent mean and standard errors of 3 independent experiments. For all data, the paired t-test was used to examine statistical significance (*p < 0.05; **p < 0.01; ***p < 0.005; n.s., non-significant).

### In MTCL2-depleted CGNs, dendrite extensions are biased in the direction opposite to axon elongation

While characterizing MTCL2-depleted CGNs, we observed that their dendrite extensions were biased in the opposite direction of axon elongation (see Fig. 2A,C). To confirm this tendency, we quantitatively estimated the direction of dendrite extension in the CGNs at DIV6. Angles were measured based on the positions of the base of each dendrite (MAP2-positive neurite) and an axon (MAP2-negative, thin neurite) on the cell body (see Fig. 3A for an illustration). Consistent with previous reports (Powell et al., 1997; Zmuda and Rivas, 1998), control cells extended dendrites evenly in all directions (Fig. 3B). Conversely, dendrite extension in MTCL2-depleted CGNs was weakly or strongly biased toward the same or opposite directions of axon elongation, respectively, when focusing on dendrites longer than 15 μm (Fig. 3B). The directional bias (r), defined as the ratio of the number of dendrites extending forward or backward versus left or right, was 1.22 ± 0.05 and 1.66 ± 0.10 for control cells and MTCL2-depleted cells, respectively (Fig. 3C). However, expressing RNAi-resistant MTCL2 in MTCL2-depleted cells suppressed the directional bias of dendritic extension to 1.01 ± 0.03 (Fig. 3B,C). These results suggest that MTCL2-depleted CGNs exhibit bipolarity in relation to the direction of axon elongation, extending fewer and longer dendrites in that direction (Fig. S3E).

**Figure 3.**
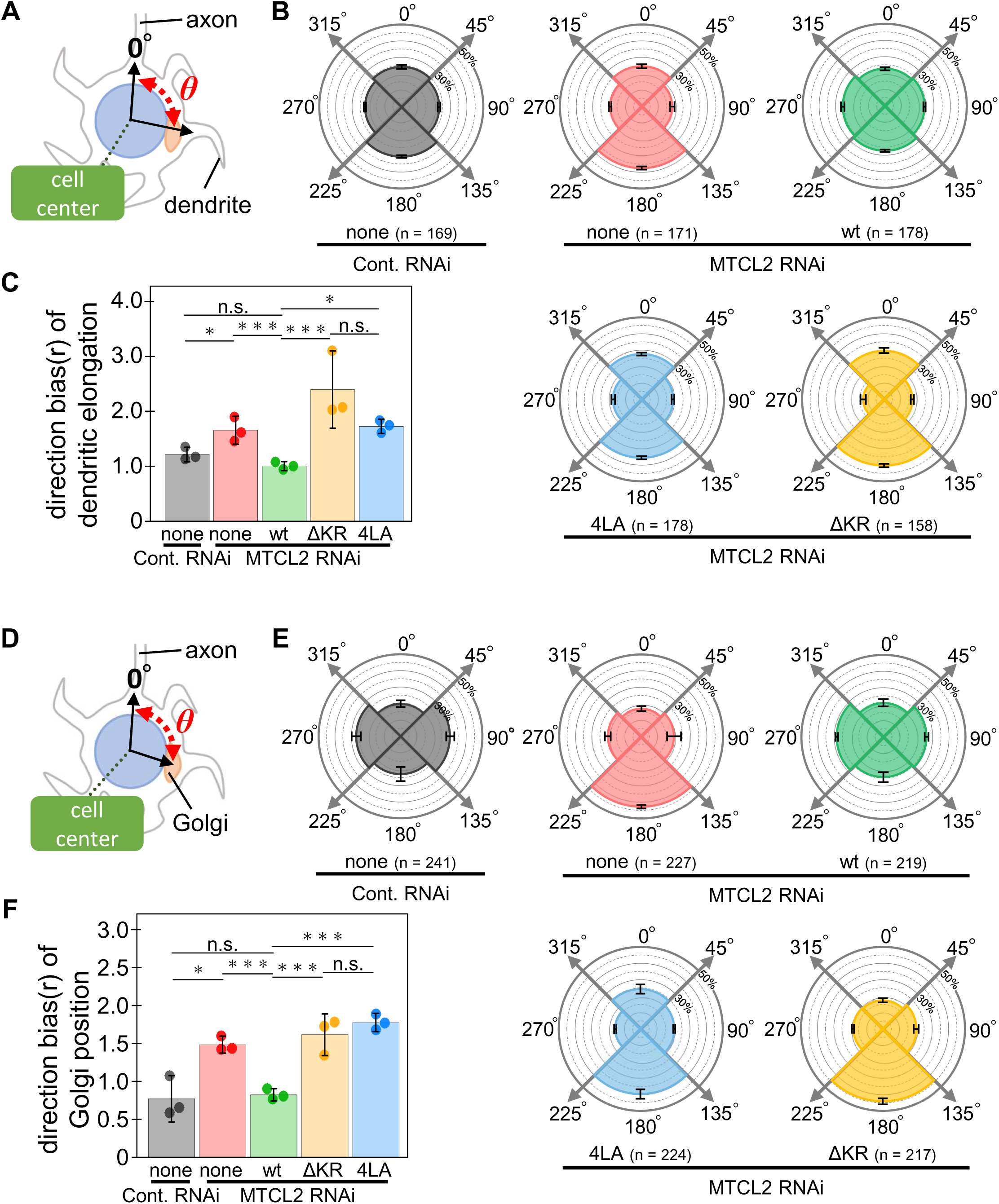
Dendrite extensions and Golgi apparatus positions of MTCL2-depleted CGNs biased in the direction opposite to axonal elongation. (A) The extension directions of dendrites longer than 15 μm were defined as the angle (*θ*) formed between the roots of the dendrite and an axon, as depicted. (B) *θ* distribution of individual dendrites of CGNs subjected to the indicated RNAi/rescue conditions and fixed at DIV6. The *θ* values were categorized in 4 directions and presented in polar histograms as indicated. More than 350 dendrites from the indicated number (n) of cells from 3 independent experiments were analyzed for each condition. (C) Comparison of dendritic direction bias (r) between the conditions. r =1 indicates no direction bias. (D) The position of the Golgi apparatus for individual CGNs was estimated as the angle (*θ*) formed between the center of the Golgi apparatus and the roots of an axon, as depicted. (E) *θ* distribution of the Golgi position in CGNs subjected to the indicated RNAi/rescue conditions and fixed at DIV6. The *θ* values were categorized in 4 directions and presented in polar histograms as indicated. The numbers (n) of cells from 3 independent experiments were analyzed. (F) Comparison of direction bias (r) of the Golgi position under each condition. Mean values and standard errors are shown for all quantitative data for each condition. In (C) and (F), the paired t-test was used to examine statistical significance (*p < 0.05; ***p < 0.005; n.s., non-significant).

### MTCL2 is necessary for the transition of dendrite extensions from bipolar to multipolar

Many studies have examined dendrite growth in CGNs using *in vitro* culture systems (Huang et al., 2007; Litterman et al., 2011; Tan et al., 2013; Kawaji et al., 2004). However, to our knowledge, no study has examined the polarity of dendrite extensions during CGN differentiation. Thus, we next examined whether the bipolar dendrite extensions observed in MTCL2-depleted cells corresponded to an intermediate state of normal CGN differentiation or an aberrant state of the cells that is not observed in normal CGN differentiation. To this end, we assessed the polarity of CGN dendrite extensions at the middle stage (DIV3) when MAP2-positive dendrites begin to appear (Fig. 1D; Fig. 4). Immunostaining with the axon marker Ankyrin G and the dendrite marker MAP2 revealed that all of the CGNs that we analyzed had a single, long axon and multiple dendrites (an average of 3.4 dendrites per cell) (Fig. 4A). This indicates that these cells have exited the initial bipolar stage with two axons and entered a stage in which they elongate multiple dendrites. In these DIV3 CGNs, the directions of the dendrite extensions were strongly biased toward the same axis as the axon elongation, as was observed in MTCL2-depleted cells (Fig. 4A,B; Fig. S4A). Interestingly, DIV3 CGN dendrites were fewer but longer than DIV5 CGN dendrites (Fig. 4C; Fig. S4B). This also corresponded to the situation observed in MTCL2-depleted cells. These results support the notion that MTCL2 knockdown arrests the bipolar-to-multipolar transition of dendrite extension, which normally occurs during CGN differentiation.

**Figure 4.**
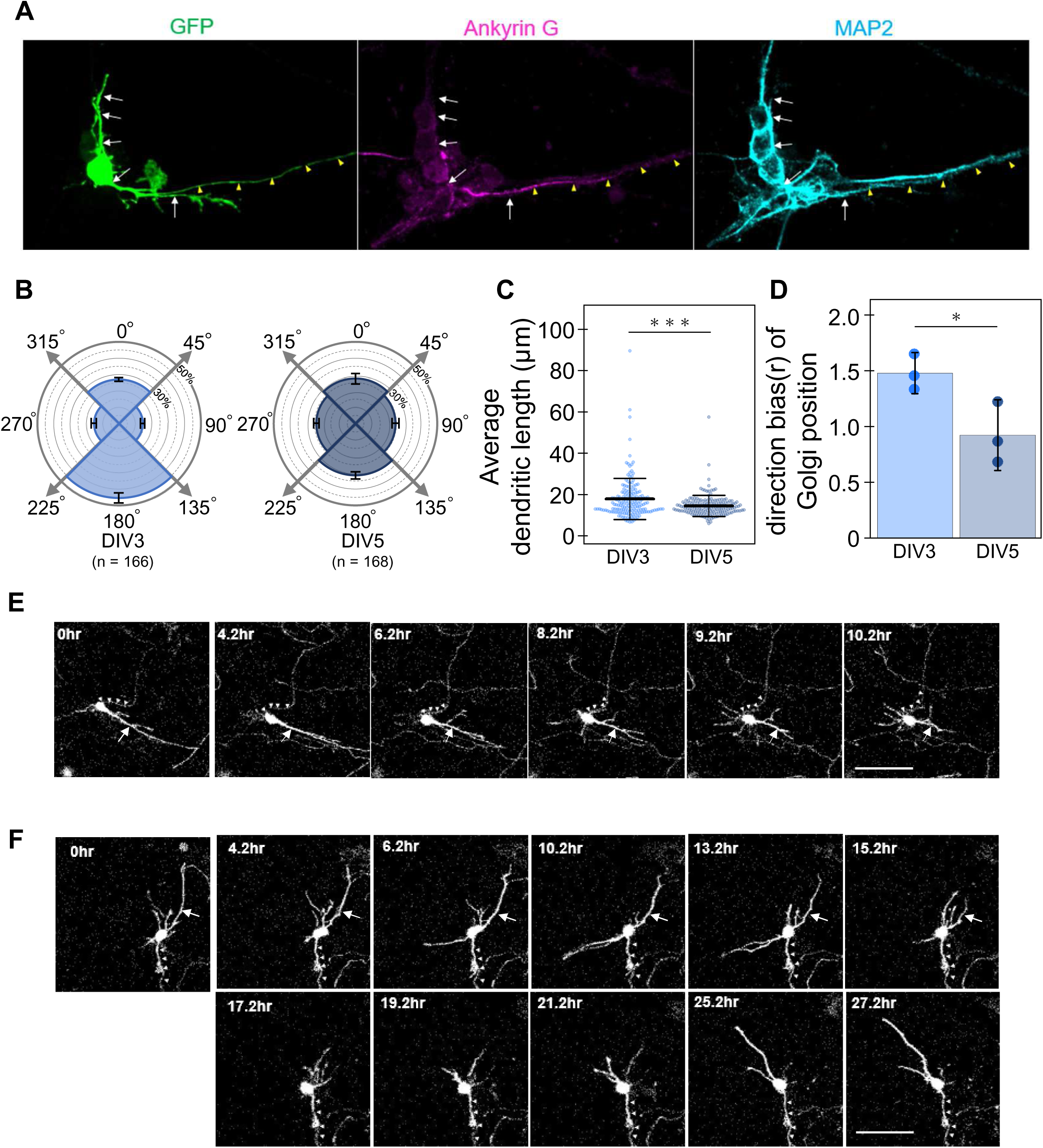
MTCL2 is required for the bipolar-to-multipolar dendrite extension transition. (A) Representative morphologies of CGNs at DIV3 visualized by GFP. Axons and dendrites were identified by staining with Ankyrin G and MAP2, respectively. Arrows and arrowheads indicate dendrites and an axon of the GFP-expressing cell, respectively. Scale bar: 20 μm. (B) *θ* distribution of individual dendrites of CGNs at DIV3 and 5. Data are presented as polar histograms for >350 dendrites of the indicated number (n) of cells from 3 independent experiments for each condition, as in Fig. 3B. (C) Distribution of the average dendritic length of individual CGNs analyzed in (B). Each data point represents the result of one neuron in 3 independent experiments. Mean values and standard deviations are shown for each scatter plot. (D) Comparison of the direction bias (r) of the Golgi position in the GCNs at DIV3 and DIV5. The mean values and standard errors of 3 independent experiments are shown for each condition. In (C) and (D), the paired t-test was used to examine statistical significance (*p < 0.05; ***p < 0.005). (E, F) Representative live imaging data of GFP-expressing CGNs transfected with control (E) or MTCL2 (F) knockdown vectors. In each case, bipolar CGNs were selected at DIV3, and recordings were performed for 48 h. The frames showing the characteristic changes in the dendrite extensions are only presented for each CGN (for comprehensive images, refer to Movies 1 and 2). Arrowheads indicate axons. Arrows indicate initial stable dendrites. Note that the MTCL2-depleted CGN exhibited shrinkage of the preexisting bipolar dendrites but failed to persistently extend short dendrites in multiple directions. Instead, the cell stably extended long bipolar dendrites, again. Scale bars: 50 μm.

To directly examine this notion, we subjected GFP-expressing CGNs, with or without MTCL2 knockdown, to live imaging at DIV3. We recorded the morphological changes every 2 min for 48 h (Fig. 4E,F; Movie 1 and 2). Consistent with the aforementioned results, the cells exhibited extended bipolar neurites and a thin, long axon (arrowheads) at the beginning of the recording (Fig. 4E,F). The control cells exhibited a thick neurite extending in the direction opposite to the axon (arrow in Fig. 4E), which remained relatively stable for a few hours. In addition, several short neurites grew and shrank dynamically in the same direction as axon elongation. A few hours after recording began, the thick neurite started shrinking (6.2-9.2 h). Simultaneously, short dynamic neurites began to elongate from various sites on the cell body (after 8.2h), and the cell eventually acquired a multipolar morphology with short dendrites (Fig. 4E; Fig. S4D; Movie 1). At this stage, the initial thick neurite became indistinguishable from the other neurites in terms of length, thickness, and dynamic nature. These results support the notion that CGN dendrites undergo a bipolar-to-multipolar transition during their differentiation. Similar shrinkage of an initial bipolar neurite (arrow) (13.2-17.2 h) and *de novo* elongation of unstable short dendrites (15.2-21.2 h) were observed in MTCL2-depleted CGNs (Fig. 4F). However, the cells could not steadily extend short dendrites longer than the width of their cell bodies (∼ 15μm) in multiple directions. Instead, these cells finally elongated new, long, stable dendrites along the initial bipolar axis (25.2 h ∼) (Fig.4F; Fig. S4D; Movie 2). These results suggest that MTCL2 is essential for disrupting the bipolarity of dendrite extension that occurs during normal CGN differentiation.

### MTCL2 promotes the bipolar-to-multipolar transition of dendrite extension through its MT- and Golgi-binding activity

The localized positions of the Golgi apparatus are closely associated with the development of various neurite processes, including axons, dendrites, and leading processes involved in neuronal migration (Horton et al., 2005; Matsuki et al., 2010; O’Dell et al., 2012; Tanabe et al., 2010; Zmuda and Rivas, 1998). Because MTCL2 is a microtubule-regulating protein that binds to the Golgi membrane (Matsuoka et al., 2020), we next assessed the possibility that MTCL2 facilitates the bipolar-to-multipolar transition of dendrite extension polarity by affecting Golgi localization. To this end, we first examined the localization of the Golgi apparatus in MTCL2-depletecd cells (Fig. 3D-F). Consistent with a previous report (Zmuda and Rivas, 1998), the Golgi apparatus did not exhibit any specific localization in DIV6 CGNs. However, in MTCL2-depleted CGNs, the Golgi position was clearly biased toward the direction opposite to axon elongation, similarly to dendrite elongation. This abnormality was randomized again by expressing RNAi-resistant MTCL2. Interestingly, CGNs at DIV3 exhibited a similarly biased Golgi localization (Fig. 4D; Fig. S4C). These results support the possibility that MTCL2 disrupts the bipolarity of dendrite extensions by affecting the Golgi position.

To evaluate this possibility, we examined the ability of MTCL2 mutants to rescue the defects exhibited by MTCL2-depleted CGNs (Fig. 2; Fig. 3; Fig. S3). Unlike wild-type MTCL2, MTCL2 mutants lacking MT-(ΔKR) or Golgi-(4LA) binding activity (Matsuoka et al., 2022) failed to rescue the biased polarity of dendrite elongation and the Golgi localization (Fig. 3B,C and E,F). In addition, they did not restore the reduced number of dendrites observed in MTCL2-depleted CGNs (Fig. 2D; Fig.S3B). The only exception was that the 4LA mutant, but not the ΔKR mutant, exhibited substantial rescue activity for dendrite length (Fig. 2E,F; Fig. S3C,D). Consequently, MTCL2-depleted CGNs expressing RNAi-resistant MTCL2 4LA developed a few short dendrites with biased polarity (Fig. S3E). This indicates that the sustained destabilization of a bipolar long neurite, one of the two events observed in the bipolar-to-multipolar transition of dendrite extensions (Fig.4E; Movie 1), does not require MTCL2’s Golgi-binding activity. In any case, these results suggest that the MT- and Golgi-binding activities of MTCL2 are crucial for disrupting the preexisting bipolarity and inducing multidirectional dendrite development in CGNs.

### During the late stage of CGN polarization, the Golgi apparatus exhibited upward extension along the microtubule cage surrounding the nucleus

To investigate how MTCL2 promotes the bipolar-to-multipolar transition of dendrite extension, we next examined the subcellular localization of MTCL2 in greater detail in relation to MTs and the Golgi apparatus.

Consistent with our previous observations in HeLa-K cells, basal confocal images of mature CGNs (DIV5) revealed intermittent granular MTCL2 stainings on MT bundles passing through MAP2-positive dendrites (Fig. S5A). In addition, numerous MTCL2 signals were observed in the cell bodies (Fig. 5A; Fig. S5A). These signals frequently colocalized with an MT network in the cell bodies that was efficiently stained by anti-tyrosinated but not anti-acetylated tubulin antibodies (Fig. S5B,C). Serial analysis of z-stack confocal images (Fig. 5A,B) and their reconstructed z-sectional views (Fig. 5C; Movie 3) revealed that the MT network completely encircles the nucleus. This indicates that it corresponds to the MT cage, the presence of which has been reported in CGNs (Rivas and Hatten, 1995; Solecki et al., 2004; Umeshima et al., 2007; Xie et al., 2003). Preferential staining with anti-tyrosinated, but not anti-acetylated, tubulin antibodies corroborated this conclusion (Umeshima et al., 2007).

**Figure 5.**
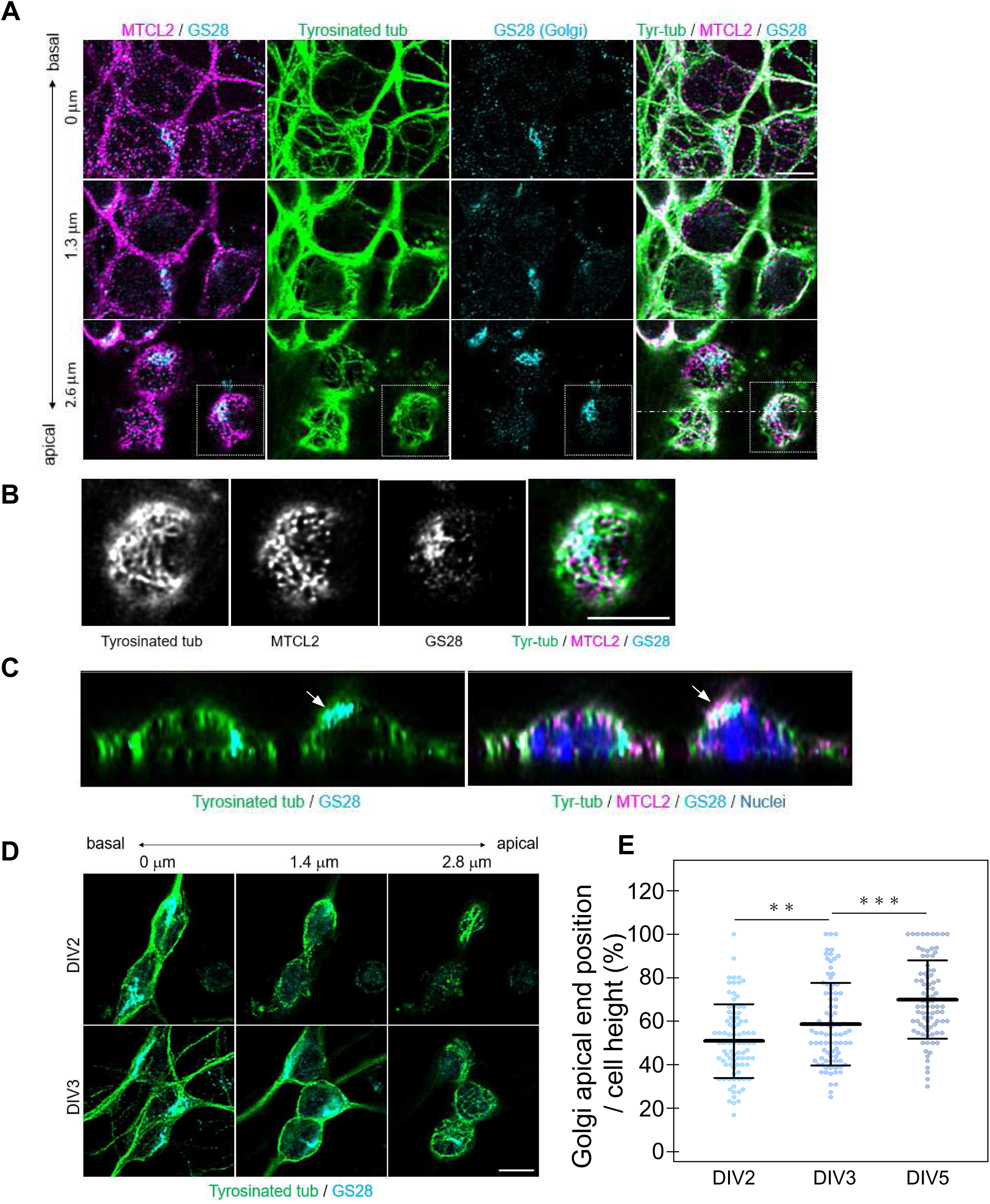
The Golgi apparatus in the differentiated CGNs was sequestered into the MT cage surrounding the nucleus. (A) High-resolution confocal images of mature CGNs (DIV5) stained for tyrosinated tubulin, MTCL2, and GS28 (Golgi membrane protein). Three images of the same field in different confocal planes are presented from the basal to apical planes, as indicated. Scale bar: 5 μm. (B) Boxed regions in (A) were enlarged to show substantial colocalization of tyrosinated tubulin, MTCL2, and the Golgi apparatus. Scale bar: 5 μm. (C) Cross-sectional views of the z-stack images along the dotted line in (A). Two types of merged images with different antibody combinations are presented. The arrows indicate the Golgi apparatus at the top of the nucleus. Scale bar: 2 μm. (D) High-resolution confocal images of immature CGNs at DIV2 and DIV3 after staining with tyrosinated tubulin and GS28. Scale bar: 5 μm. (E) Height distribution of the apical edge of the Golgi apparatus estimated as a percentage of the cell heights (see Fig. S7). Each data point represents the result of one neuron observed in 3 independent experiments. Mean values and standard deviations are shown for each scatter plot. The paired t-test was used to examine statistical significance (**p < 0.01; ***p < 0.005).

Interestingly, the Golgi apparatus in CGNs at DIV5 is often associated with the MT cage on the lateral or apical surfaces of the nucleus (Fig. 5A,B). Consequently, the Golgi apparatus was drawn toward the nucleus, extending its apical end toward the upward direction along the nuclear surface (Fig. 5C). In the most extreme cases, the entire apparatus was placed on top of the nucleus (see arrows in Fig. 5C; Movie 3). This differs greatly from the situation observed in CGNs at DIV2. At this stage, the Golgi apparatus was found in the basal position, biased toward the base of one of the bipolar processes, as previously reported (Fig. 5D; Fig. S6A,B) (Zmuda and Rivas, 1998). Some DIV2 cells contained cortical MTs underneath the apical membranes of their cell bodies (Fig. 5D, see the arrowhead in Fig. S6B). However, these MTs were not organized into a dense, isotropic network. Rather, they tended to align parallel to the preexisting bipolar axis (Fig. 5D). Even at the CGNs at DIV3, where isotropic network-like MT structures were detected (Fig. 5D, Fig. S6A), the upward shift of the Golgi apparatus was not evident (Fig. 5D; Fig. S6B). Quantitative analysis demonstrated that the cell heights of the CGNs at DIV3 and DIV5 did not differ significantly (Fig. S7A,B). However, the height of the apical end of the Golgi apparatus relative to the corresponding cell height increased as the polarization proceeded (Fig. 5E). These results suggest that the final CGN polarization is tightly coupled with the sequestration of the Golgi apparatus from the base of the bipolar neurites into an MT cage around the nucleus. This is consistent with a previous report that the Golgi apparatus in mature CGNs is not localized to the base of any neurites, including axons and dendrites (Zmuda and Rivas, 1998)

### MTCL2 promotes the sequestration of the Golgi apparatus into the MT cage

Importantly, in the CGNs at DIV5, many MTCL2 signals substantially overlapped with those of the Golgi and MT signals around the nucleus (Fig. 5A-C). Since both the MT- and the Golgi-binding activities of MTCL2 were necessary to disrupt the Golgi’s biased localization in the x-y plane (Fig. 3E,F), these results suggest that MTCL2 plays a crucial role in the Golgi’s upward extension by facilitating its association with the MT cage. To verify this possibility, we reexamined the positions of the Golgi apparatus in MTCL2-depleted cells in the z direction (Fig. 6). As expected, the positions of the Golgi apparatus were significantly lowered in MTCL2-depleted cells (Fig. 6A). Quantitative analysis demonstrated that MTCL2 knockdown did not change the cell height (Fig. S7C). However, the apical ends of the Golgi apparatus were lowered to a level similar to that observed in bipolar CGNs at DIV3 (Fig. 6C). This defect was restored by expressing RNAi-resistant MTCL2 wild type but not by expressing MTCL2 ΔKR or 4LA (Fig. 6C). These results support the idea that MTCL2 extends the Golgi apparatus upward along the nucleus by connecting it to the MT cage (Fig. 7A).

**Figure 6.**
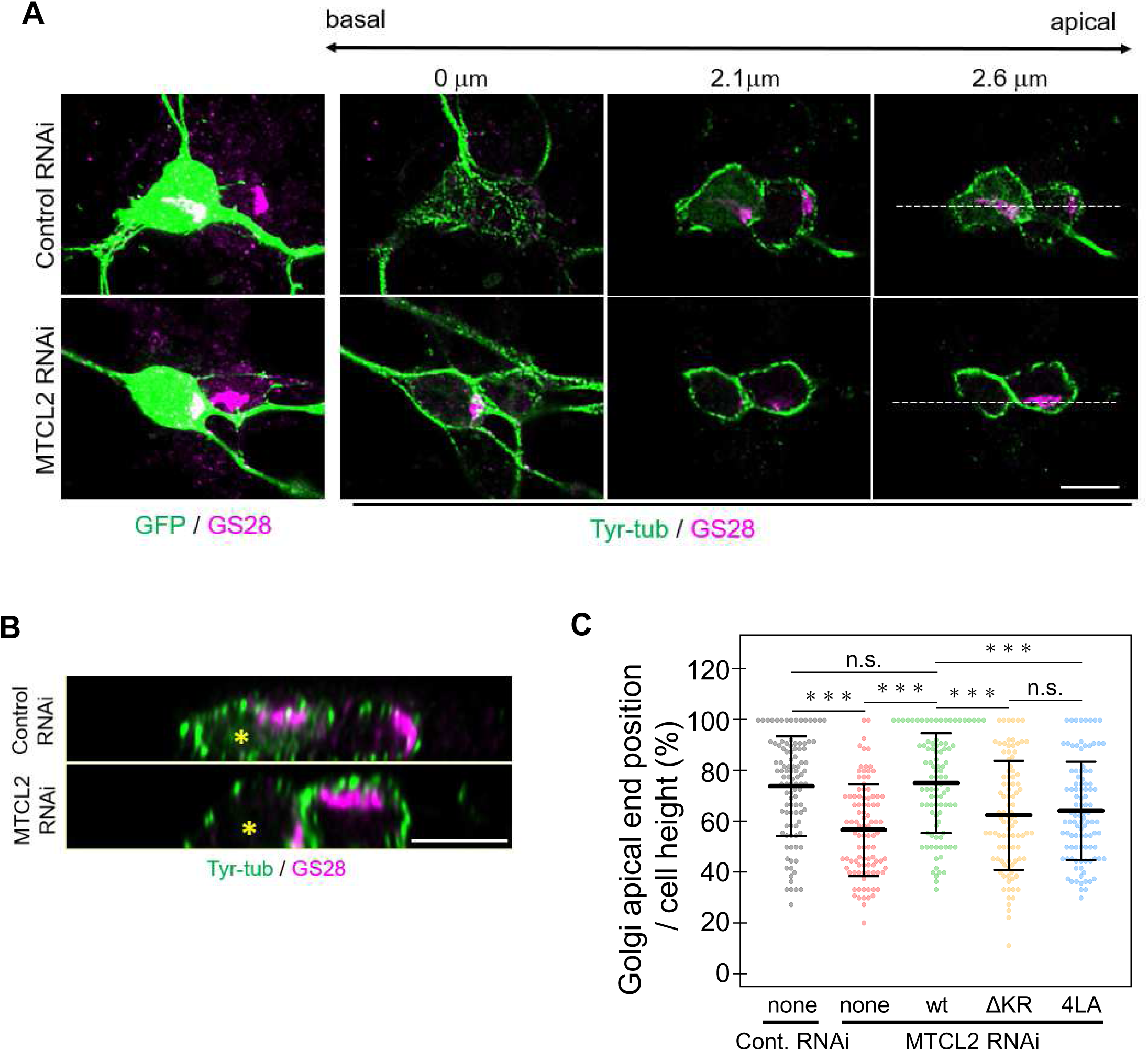
MTCL2 promotes the upward extension of the Golgi apparatus in an MT- and Golgi-binding activity-dependent manner. (A) High-resolution confocal images of CGNs subjected to the indicated RNAi, fixed at DIV6, and stained for tyrosinated tubulin and GS28. On the left, projected views of the z-stack images indicate cells expressing the indicated shRNA vector together with GFP. Images obtained at three different confocal planes are shown on the right. Scale bar: 5 μm. (B) Cross-sectional views of the z-stack images along the dotted lines in (A). Asterisks indicate GFP-positive cells. Scale bar: 5 μm. (C) Height distribution of the Golgi apical edge in CGNs subjected to the indicated RNAi/rescue conditions and fixed at DIV6. Each data point represents the result of one neuron observed in 3 independent experiments. Mean values and standard deviations are shown for each scatter plot. The paired t-test was used to examine statistical significance (***p < 0.005; n.s., non-significant).

**Figure 7.**
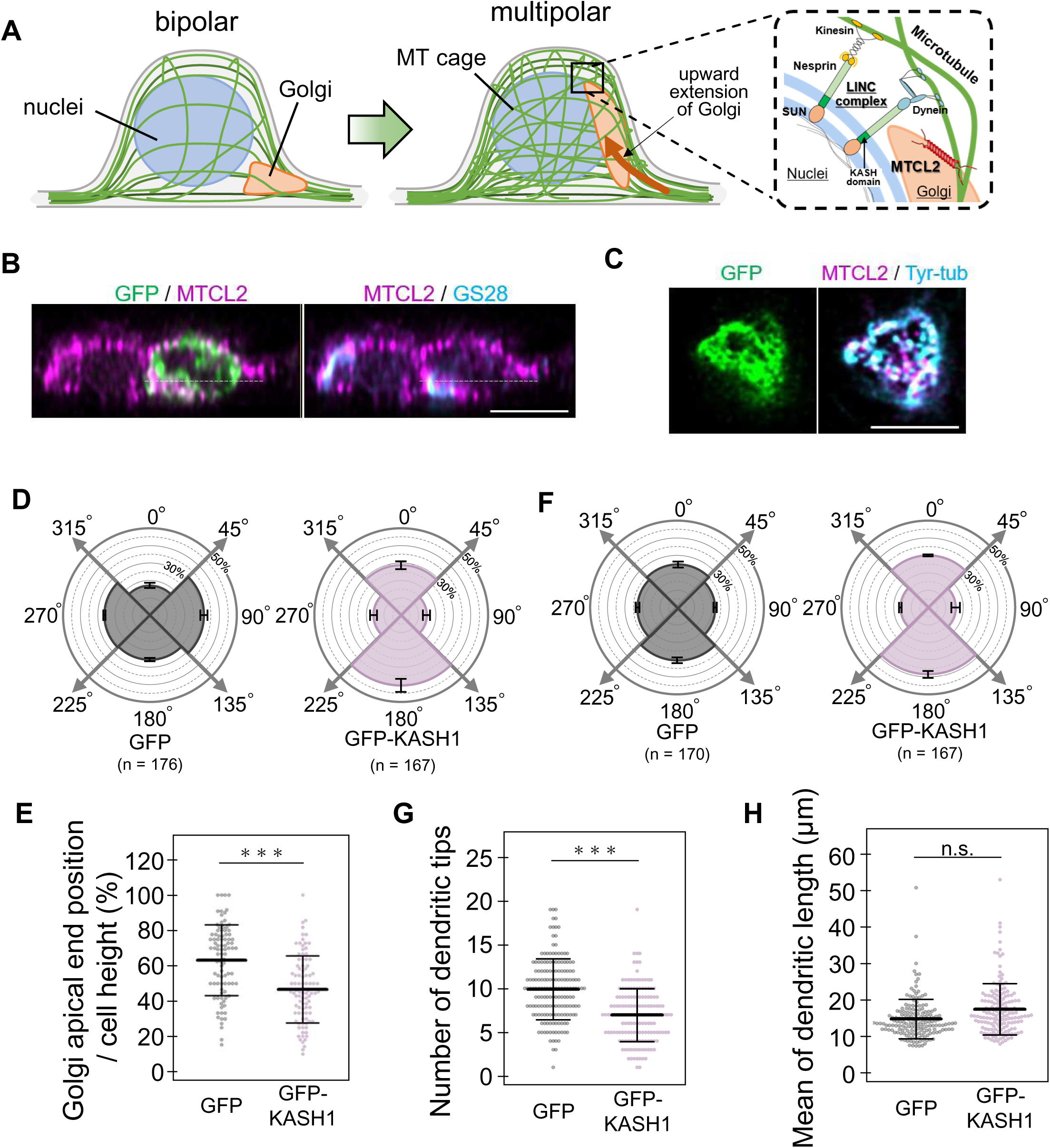
Inhibition of LINC complex activity mimicked most of the phenotypes shown by MTCL2 knockdown. (A) Schematic illustration of our working hypothesis. At the late phase of CGN polarization, the MTCL2-mediated association between the Golgi apparatus and the MT cage induces an upward extension of the Golgi apparatus along the nucleus. This recruitment to the MT cage results in sequestration of the Golgi apparatus from the roots of the preexisting bipolar dendrites. The molecular organization of the LINC complex on the nuclear membrane is illustrated on the right. (B) A typical cross-sectional view of CGNs expressing GFP-KASH1, fixed at DIV6, and stained for MTCL2 and GS28. The white dotted lines indicate the focus plane from which the confocal image shown in Fig. S8A was obtained. Scale bar: 5 μm. (C) An apical confocal section of a GFP-KASH1-expressing cell stained for MTCL2 and tyrosinated tubulin. Scale bar: 5 μm. (D) *θ* distribution of the Golgi position in CGNs expressing GFP or GFP-KASH1, examined at DIV6. Data are presented as polar histograms as shown in Fig. 3E for the indicated number (n) of cells from 3 independent experiments. (E) Hight distribution of the Golgi apical edge in the same CGNs analyzed in (C). (F) *θ* distribution of individual dendrites of CGNs expressing GFP or GFP-KASH1 and examined at DIV6. Data are presented as polar histograms as shown in Fig. 3B for >400 dendrites of the indicated number (n) of cells from 3 independent experiments. (G, H) Distribution of the number of dendritic tips (G) and averaged dendritic length (H) of the same CGNs analyzed in (F). Each data point in (E), (G) and (H) represents the result of a single neuron. Mean values and standard deviations are shown for each scatter plot. The paired t-test was used to examine statistical significance (***p < 0.005; n.s., non-significant).

### The LINC complex activity is necessary for sequestering the Golgi apparatus into the MT cage

The initial observation of the MT cage in CGNs was 30 years ago (Rivas and Hatten, 1995), but its molecular basis remains unclear. The LINC complex is a potential candidate associated with the MT cage because it links the nuclear membrane to MTs (Chang et al., 2015; Kengaku, 2018). Recent studies have revealed the indispensable role of the LINC complex in nuclear rotation, deformation, and translocation during CGN migration (Wu et al., 2018; Zhou et al., 2024). Thus, we investigated the role of the LINC complex in the bipolar-to-multipolar transition of dendrite elongation to confirm the involvement of the MT cage in this process.

The Nesprin protein is a component of the LINC complex located in the outer nuclear membrane. It interacts with MT motors via its N-terminal cytoplasmic stretch. Another component is the inner nuclear membrane protein SUN, which interacts with the nuclear lamina and chromatin within nucleus via its N-terminus (Fig. 7A). Nesprins interact with SUN proteins through their C-terminal KASH domains in the perinuclear space (Fig. 7A). Consequently, the overexpression of a Nesprin-1 mutant lacking the cytoplasmic stretch (KASH1) inhibits LINC complex activity by dominantly interfering with the interaction between endogenous Nesprin-1/2 and SUN proteins (Stewart-Hutchinson et al., 2008; Wu et al., 2018). When expressed in primary cultured CGNs, GFP-KASH1 localized to the nuclear membrane, as expected (Fig. 7B). Close inspection revealed a network-like distribution of GFP-KASH1 on the nuclear membrane, likely reflecting the underlying nuclear lamina pattern via SUN proteins (Fig. 7C; Fig. S8A). These cells exhibited the formation of the MT cage itself (Fig. 7C; Fig. S8B) and the colocalization of MTCL2 with the Golgi apparatus and the MT cage (Fig. 7B,C; Fig. S8A). However, the randomization of the Golgi position in the x-y plane (Fig. 7D; Fig. S8C) and the upward extension of the Golgi apparatus in the z direction (Fig. 7B,E) were specifically suppressed in these cells, as observed in MTCL2-depleted CGNs.

Importantly, when we examined the polarity of dendrite extensions, cells expressing GFP-KASH1 exhibited a reduced number of dendrites with a biased extension direction to the preexisting bipolar axis (Fig. 7F,G; Fig. S8D). As these cells did not exhibit a significant increase in dendritic length (Fig. 7H; Fig. S8E), they perfectly mimicked the phenotypes exhibited by MTCL2-depleted CGNs expressing the MTCL2 4LA mutant, which lacks Golgi-binding activity (Fig. S3E, Fig. S8F). These results support the hypothesis that the MTCL2-mediated sequestration of the Golgi apparatus within the MT cage plays a pivotal role in the multipolar development of dendrites observed in CGNs during the final stage of their polarization.

## Discussion

Previously, we reported that MTCL2 is a unique MT-regulating protein with Golgi-binding activity (Matsuoka et al., 2022). In this study, we demonstrated that MTCL2 is highly expressed in CGNs and plays an indispensable role in the later stage of polarization, during which the dendrite extensions undergo a bipolar-to-multipolar transition. We provided data indicating that MTCL2 facilitates this transition by mediating the sequestration of the Golgi apparatus into the MT cage surrounding the nucleus.

The unique polarization process observed in isolated CGN cultures was first documented in 1997 (Powell et al., 1997;Kollins et al., 1999; Zmuda and Rivas, 1998). These pioneering studies also demonstrated the significance of the Golgi apparatus’s positioning in the development of initial bipolar axons (Zmuda and Rivas, 1998). However, the mechanism by which the initial bipolarity is disrupted to result in multipolar dendrite extension remains to be elucidated. In this paper, to understand the molecular basis of the MTCL2 function in CGNs, we focused on this issue and demonstrated that the initial extension of multiple dendrites exhibits bipolarity (Fig. 4A, B). The live imaging analysis of CGNs from DIV3 to DIV5 not only corroborated this conclusion but also revealed that this bipolarity is disrupted through two distinct processes: destabilization of the stable dendrites elongating opposite to an axon, and the *de novo* generation of unstable dendrites from the multiple sites of the cell body (Fig. 4E,F). We further found that MTCL2 knockdown impeded both processes, resulting in a reduced number of longer dendrites that exhibit an extension bias directed in the opposite direction of an axon (Fig. 3A-C). The rescue experiments further indicated that the MT-binding activity of MTCL2 was required for both processes, whereas the Golgi-binding activity was dispensable for the sustained destabilization of a bipolar long neurite and only required for the persistent generation of multipolar dendrites from the cell body. Actual CGNs *in vivo* undergo a second bipolar phase between the initial bipolar phase (characterized by two axons) and the final complex morphology (characterized by multipolar dendrites). During the second bipolar phase when CGNs migrate deep into the internal granular layer along the radially aligned Bergmann glial cells, by extending a leading process in the migration direction and drawing the axon that elongates to the opposite direction (trailing process) (Gao and Hatten, 1993; Solecki et al., 2006). Importantly, Kawaji et al. (2004) demonstrated that the leading process observed in this phase exhibits dendritic properties and ultimately contracts, resulting in differentiation into one of the multipolar dendrites. Therefore, it is highly plausible that the bipolarity observed in DIV3 CGNs with single axons mimics that exhibited by migrating CGNs. Furthermore, the stable dendrites that elongate in the direction opposite to the axon likely correspond to the radial leading process of CGNs *in vivo*. Thus, the present results support the notion that MTCL2 is required to disrupt the bipolarity of CGNs established during migration.

At present, the mechanism by which the MT-binding activity of MTCL2 is required for the destabilization of long bipolar dendrites observed at DIV3 remains unclear. Further research is necessary to elucidate this issue. However, the present study provides data indicating how the MT- and Golgi-binding activities of MTCL2 promote the multipolar development of *de novo* dendrites. Our novel findings, which include the observation of upward extension of the Golgi apparatus of CGNs in the later stage of polarization along the nucleus in an MTCL2-dependent manner (Fig. 5), support the proposal that MTCL2 facilitates the bipolar-to-multipolar transition of dendrite extension by mediating the association between the Golgi apparatus and the MT cage. This, in turn, promotes the sequestering of the Golgi apparatus to the MT cage from the base of preexisting bipolar dendrites, thus inhibiting the polarized vesicle transport required for polarized dendrite extension (Fig. 7A). This hypothesis was further substantiated by the finding that the inhibition of the normal formation of the LINC complex also resulted in the suppression of the bipolar-to-multipolar transition of the dendritic extension. Notably, the inhibition of the LINC complex did not affect the formation of the MT-cage; rather, it suppressed the upward extension of the Golgi apparatus. The results indicate that the pulling forces generated by the MT motors may contribute to the attraction of the Golgi apparatus to the MT cage, thereby inducing its upward extension (Fig.7A).

The critical role of the Golgi apparatus in dendrite development has been extensively studied (Horton et al., 2005; Nakagawa and Iwasato, 2023; Tanabe et al., 2010; Wu et al., 2015). However, most of these studies have focused on the analysis of pyramidal neurons located in the cerebral cortex and hippocampus, as well as on Purkinje cells in the cerebellum. These neurons all develop thick, apically polarized primary dendrites, also known as apical dendrites. These studies demonstrate that polarized Golgi localization is required for asymmetric dendrite growth. In contrast, the present study distinguished itself by analyzing CGNs that exhibit near-symmetric short dendrites with uniform length, and demonstrated that sequestering the Golgi apparatus from the base of any dendrite is essential for developing multipolar dendrites devoid of any asymmetry. Furthermore, during this analysis, the MT cage was identified as the target structure to which the Golgi apparatus was sequestered. Since the MT cage had previously been studied only in terms of its role in nucleokinesis observed in neural cell migration (Rivas and Hatten, 1995; Solecki et al., 2004; Xie et al., 2003), the present finding reveals an unexpected function of this higher-order MT structure. The molecular mechanism suggested in this study may be specific to CGNs. Indeed, the presence of the MT cage has primarily been described in CGNs, but not reported conspicuously in other neurons (Rivas and Hatten, 1995; Solecki et al., 2004; Umeshima et al., 2007). However, many nonpyramidal neurons such as GABAergic interneurons lack apical dendrites and develop symmetrical, uniform dendrites (Horton et al., 2005; Müller et al., 1984). In addition, some pyramidal neurons remodel their asymmetric dendrites by withdrawing their apical dendrites and acquiring a multipolar morphology (Koester and O’Leary, 1992). Therefore, neurons other than CGNs may utilize the MT-cage-dependent regulation of the Golgi position, at least in part.

In summary, the results of this study shed light on how CGNs extend dendrites in multiple directions. Our results highlight the significance of the MT cage surrounding the nucleus for removing the Golgi apparatus from the base of preexisting neurites. The critical importance of the MTCL2-mediated association between the Golgi apparatus and the MT cage, as well as that of the LINC complex-mediated fixation of MT motors to the nuclear membrane, was also suggested.

## Materials and Methods

### Animals

All protocols were approved by and performed in accordance with the guidelines established by the Institutional Animal Care and Use Committee of Yokohama City University, Medical Life Science (Approval No. T-A-19-006 and T-A-22-007).

### Antibodies

The primary antibodies used in this study were as follows: anti-SOGA1 to detect MTCL2 (HPA043992), anti-tau1 (MAB3420), anti-calbindin (C9848), anti-Ankyrin G (MABN466) from Merk Sigma-Aldrich; anti-MAP2 (sc-32791), anti-α-tubulin (DM1A: sc-32293), anti-acetylated tubulin (sc-23950), anti-tyrosinated tubulin (sc-53029), anti-GFP (sc-9996) from Santa Cruz Biotechnology; anti-GM130 (610822) and anti-GS28 (611184) from BD Transduction Laboratories; anti-GAPDH (5G4) from HyTest Ltd; and anti-NeuN (MAB377) from Merck Millipore.

### Primary Neuron Culture and Transfection

Primary cerebellar granule neurons (CGNs) were isolated from P6 neonatal mice and prepared as previously described (Krämer and Minichiello, 2010). Neurons were plated on coverslips coated with poly-D-lysin (0.1 mg/mL; Merk Sigma-Aldrich or Thermo Fisher Scientific) at a density of 180,000 cells/well, 300,000 cells/well, or 360,000 cells/well (24 well plates) and cultured for 8 h in Dulbecco’s modified Eagle medium (DMEM) (Life Technologies Corporation) containing 10% fetal bovine serum, penicillin (100 U/ml), streptomycin (0.1 mg/ml), 3.5 mM glutamine, and 0.9 mM sodium pyruvate (nacalai) under controlled temperature and CO_2_ conditions (37℃, 5%). Then, cells were subjected to plasmid transfection using a CalPhos^TM^ Mammalian Transfection Kit (Takara) according to the manufacturer’s instruction for 8∼16 h, followed by incubation with the original medium supplemented with 0.01 mM Ara-C (Tokyo Chemical Industry Co., Ltd.) and 20 mM KCl until fixation.

### Plasmids

For knockdown experiments, pEB-super-GFP vectors simultaneously expressing EGFP and appropriate shRNA were first constructed (Masuda-Hirata et al., 2009). Then, the EBNA1 cDNA sequence and the associated CMV promoter were removed to reduce vector size. For mouse MTCL2 knockdown, the RNAi target sequence 5′-gacggtcagtgtaggtcttca-3′ was inserted under the H1 promoter as a hairpin oligonucleotide sequence in a tail-to-tail fashion (Satake et al., 2017). The control shRNA sequence used was 5′-cagtcgcgtttgcgactgg-3′. To generate RNAi-resistant mouse MTCL2 expression vectors, silent mutations (<u>a</u> act gt<u>a</u> <u>tca</u> gt<u>c</u> gg<u>c</u> <u>t</u>t<u>g</u> cag a) in the RNAi target sequence were introduced in the cDNAs of V5-tagged wild-type or mutant mouse MTCL2 (4LA or ΔKR) subcloned in the pCAGGS expression vector (Matsuoka et al., 2022). In rescue experiments, the shRNA expression vector for mouse MTCL2 and appropriate expression vectors for RNA-resistant mouse MTCL2 wild type or mutant MTCL2 were co-transfected into CGNs. To construct a GFP-KASH1 expression vector, a human Nesprin-1 cDNA fragment encoding the C-terminal region (8726-8799 a.a.) was amplified by RT-PCR from total RNA from HeLa-K cells and cloned into pEGFP-C1 at BglII/BamHI sites (Stewart-Hutchinson et al., 2008). The resulting GFP-KASH1 cDNA fragment was subcloned into the pCAGGS vector to change the promoter.

### Immunocytochemistry

The CGNs were fixed in 4% paraformaldehyde (PFA) at room temperature for 15 min at 37℃. After fixation and blocking, samples were incubated with appropriate primary antibodies diluted in TBST (10 mM Tris–HCl pH 7.5, 150 mM NaCl, 0.01% [v/v] Tween 20) containing 0.1% (w/v) BSA overnight at 4 °C. After washing with PBST, samples were incubated with the appropriate secondary antibodies conjugated with Alexa Fluor 488, 555, or 647 (Life Technologies Corporation) for 60 min at room temperature. Dilutions of the used antibodies are as follows: anti-SOGA1 (1/2000), anti-α-tubulin (1:1000), anti-acetylated tubulin (1:2000), anti-tyrosinated tubulin (1:2000), anti-GM130 (1:1000), anti-GS28 (1:300), anti-GFP (1:2000), anti-MAP2 (1:1000), anti-tau1 (1:1000), and anti-Ankyrin G (1:1000). All secondary antibodies were used at a 1:2000 dilution. The nuclei were counterstained with 4’, 6-diamidino-2-phenylindole (MBL, Japan) at a 1:2000 dilution in PBST during the final wash. For image acquisition, samples on coverslips were mounted onto glass slides in Prolong Diamond Antifade Mountant (Thermo Fisher Scientific).

### Immunohistochemistry

Mice under deep isoflurane anesthesia were fixed by transcardial perfusion with 4% paraformaldehyde (PFA) in 0.1 M PB (0.1 M NaH_2_PO_4_ and 0.1 M Na_2_HPO_4_, pH 7.4). Brains were removed from the skull, post-fixed for 3–5 h at room temperature, and sectioned at 50-µm thickness using a Vibratome (VT1200S; Leica Microsystems, Wetzlar, Germany). Following permeabilization with 0.1% Triton-X-100 in 0.1 M PB with gentle shaking for 30 min, cerebellar vermis slices were blocked with 10% calf serum in TBST for 30 min at room temperature. Immunofluorescence staining was performed as previously described.

### Image acquisition and processing

Most images were acquired using a Leica SP8 laser scanning confocal microscope equipped with an HC PL APO 63x/1.40 Oil 2 objective using the Hybrid Detector in photon-counting mode. To obtain super-resolution images, HyVolution2 imaging was performed on the same system using Huygens Essential software (Scientific Volume Imaging). Single confocal images are shown unless otherwise indicated. The same confocal system was used to obtain time-lapse images, except that an HC PL APO 20x/0.75 objective was used. Cells seeded onto 35-mm glass-bottom dishes and subjected to appropriate transfection were placed in a closed heat-controlled chamber (Leica) with a CO_2_ supply system (Tokken) on the microscope stage for 2 days under autofocus control. Wide-view or low-resolution images for quantification (Figs. 2, 3, and 4) were obtained using an AxioImager ZI microscope (Carl Zeiss, Oberkochen, Germany) equipped with a Plan-Apochromat 40×/0.95 objective and an Orca II CCD camera (Hamamatsu Photonics, Shizuoka, Japan). To obtain images of brain slices and to measure the heights of cells and the apical end of the Golgi apparatus, an AxioImager Z1 microscope equipped with a CSU10 disc confocal system (Yokogawa Electric Corporation, Tokyo, Japan) and a 20x/1.4 NA or 100x/1.46 NA Plan-apochromat objective was used. Images were acquired using MetaMorph software (Molecular Devices, Sunnyvale, CA) and processed using ImageJ software to obtain appropriate brightness and contrast.

### Quantification

To quantify the enrichment of MTCL2 and MAP2 in CGN neurites, immunofluorescence signal intensities of anit-MTCL2 and anti-MAP2 staining were measured in ROIs of 1 μm^2^ circles placed on anti-acetylated tubulin- or tau1-positive neurites. To analyze the morphology of CGNs, the positions of bases, tips, or branching points of neurites (axons or dendrites) were marked manually, and their lengths and numbers of tips were automatically quantified using the NeuronJ plugin for ImageJ. Essential functions of ImageJ were used to manually analyze the polarity of the dendrite extension direction, Golgi position in the x–y plane, and number of bases of dendrites. The dendritic lengths of CGNs subjected to live imaging were also analyzed manually. The cell height and position of the Golgi apical end were determined from the focal plane at which the fluorescent signals of tyrosinated tubulin and GS28 disappeared in z-stack images taken at an interval of 0.3 μm. The 3D video shown in Movie 3 was created using software (LAS X) associated with the Leica SP8 laser scanning confocal system.

### Cell extraction and western blotting

CGN extracts were prepared by adding RIPA buffer (25 mM Tris/HCl [pH 7.5], 150 mM NaCl, 1% NP40, 1% deoxycholic acid, 0.1% SDS) containing a protease inhibitor cocktail (Merk Sigma-Aldrich, P8340), followed by brief sonication and centrifugation (15,000 × *g*, 15 min). Samples were separated using SDS-PAGE and transferred to polyvinylidene fluoride membranes. Blots were incubated in blocking buffer containing 5% (w/v) dried skim milk in PBST (8.1 mM Na_2_HPO_4_.12H_2_O, 1.47 mM KH_2_PO_4_, 137 mM NaCl, 2.7 mM KCl, and 0.05% Tween 20), followed by overnight incubation with anti-SOGA1 or anti-GAPDH antibodies diluted in blocking buffer. The diluted anti-SOGA1 and anti-GAPDH antibodies were 1:1000 and 1:5000, respectively. Blots were then exposed to horseradish peroxidase (HRP)-conjugated secondary antibodies (GE Healthcare) diluted in blocking buffer for 60 min at room temperature and washed again. Blots were visualized using an Immobilon Western Chemiluminescent HRP Substrate (Millipore) or ECL western blotting detection system (GE Healthcare). Chemiluminescence was quantified using an ImageQuant LAS4000 Luminescent Image Analyzer (GE Healthcare).

## Acknowledgments

We thank Dr. Tomoko Satake for her initial support in the establishment of the primary cerebellar granule cell culture. This research was supported by KAKENHI (to A.S.) (Grant Numbers 19H03228 and 22H02621) of the Ministry of Education, Culture, Sports, Science and Technology (MEXT) of Japan.

## Author contributions

Designed research: A.S., M.M., Performed research: M.M., A.S., Analyzed data: M.M. A.S, Wrote the paper M.M., A.S., Supervision: A.S.; Funding acquisition: A.S.

## Movie Legends

**Movie 1. Morphological changes in the control CGN from DIV3 to DIV5.**

GFP fluorescence images were taken every 2 min for 40 h. The video speed was 30 fps.

The representative frames of this movie are shown in Fig. 4E.

**Movie 2. Morphological changes in the MTCL2-depleted CGN from DIV3 to DIV5.**

GFP fluorescence images were taken every 2 min for 40 h. The video speed was 30 fps.

The representative frames of this movie are shown in Fig. 4F.

**Movie 3. 3D reconstructed images of polarized CGNs at DIV5.**

The z-stack images of the CGNs shown in Fig. 5A were reconstructed into 3D stereoscopic images with different combinations of staining.

