## Supplementary figures for "The bipolar-to-multipolar transition of dendrite extension of cerebellar granule neurons requires MTCL2 to link the Golgi apparatus to the microtubule cage"

### Figure S1

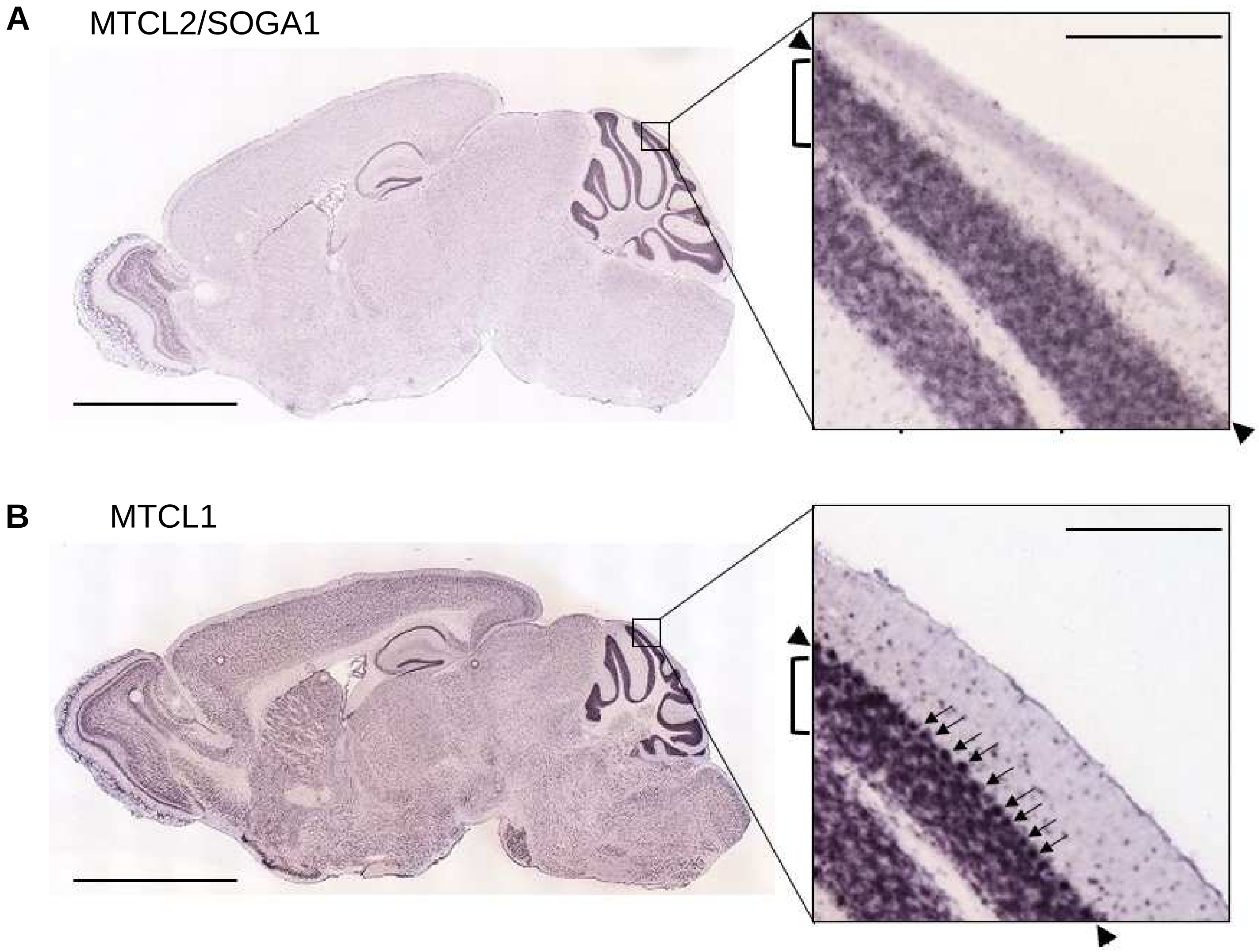


**Figure S1. MTCL2 mRNA is highly expressed in cerebellar granule neurons**

(A) MTCL2/SOGA1 mRNA expression in the adult mouse brain (age 56 weeks, male). From the Allen Mouse Brain Atlas (http://mouse.brain-map.org/experiment/show/70228227). Section 14. Arrowheads and left square brackets in the enlarged image (right panel) indicate the positions of the Purkinje cell and molecular layers, respectively. (B) Expression of MTCL1 mRNA in adult mouse brain(age 56 weeks, male). From the Allen Mouse Brain Atlas (https://mouse.brain-map.org/experiment/show/71808705). Section 13. Arrows in the enlarged image (right panel) indicate the positions of Purkinje cells. Note that MTCL1, but not MTCL2, was detected in Purkinje cells. Scale bars: 3356 μm (left) and 209 μm (right insets).

### Figure S2

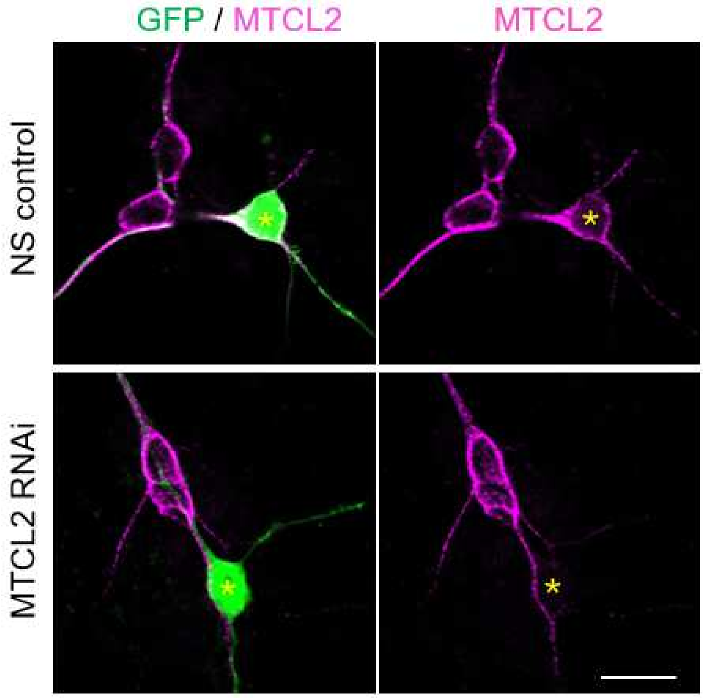


**Figure S2. MTCL2 knockdown efficiency of the shRNA expression vector**

CGNs transfected with the indicated shRNA vector were fixed at DIV6 and stained for MTCL2. The projected views of the z-stack images. Asterisks indicate GFP-positive cells that accepted the indicated shRNA vector. Scale bar : 10 μm.

### Figure S3

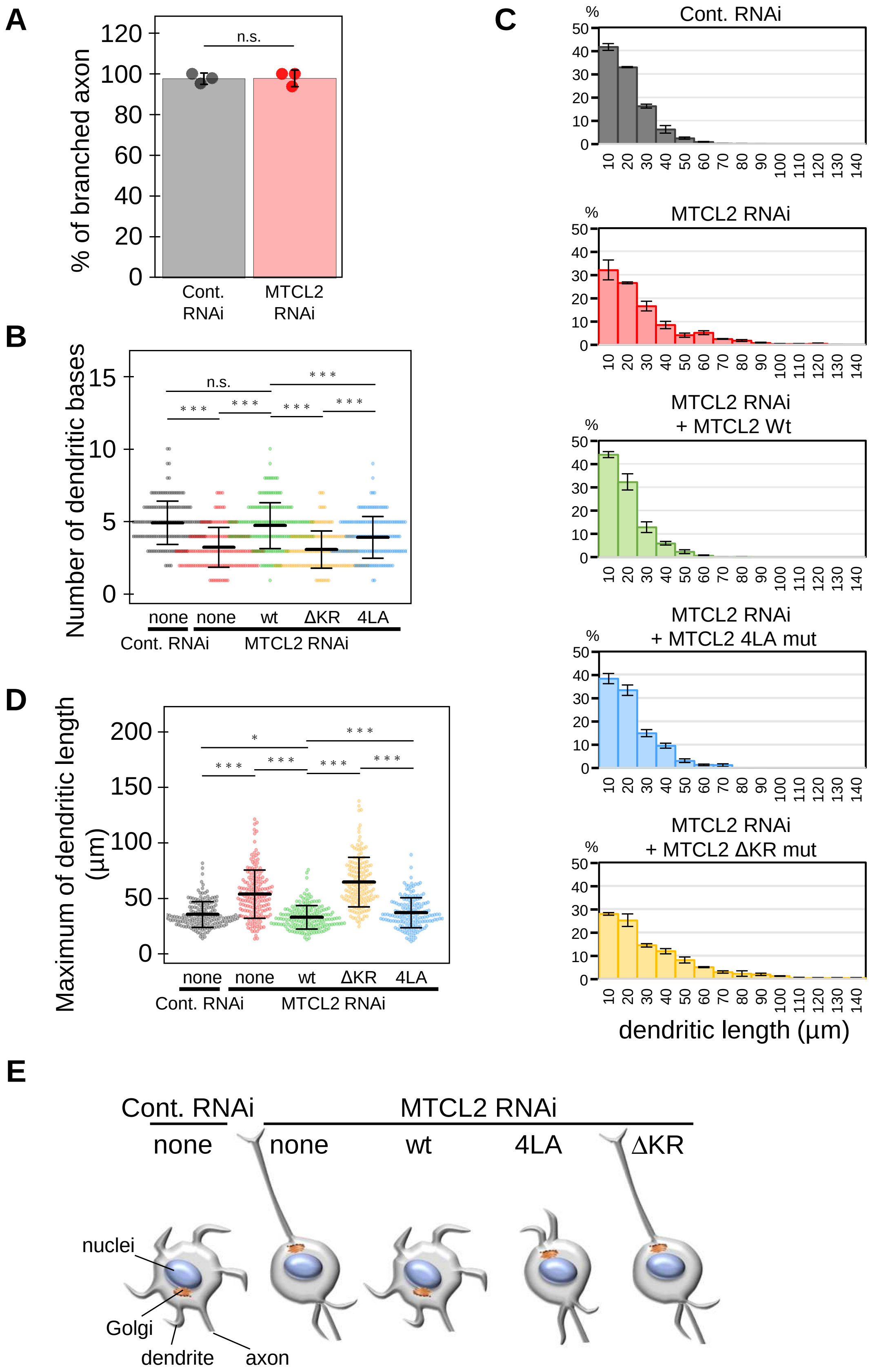
A
% of branched axon
n.s.
Cont. RNAi
MTCL2 RNAi
B
Number of dendritic bases
n.s.
* * *
none none wt ΔKR 4LA
Cont. RNAi MTCL2 RNAi
C
Cont. RNAi
MTCL2 RNAi
MTCL2 RNAi + MTCL2 Wt
MTCL2 RNAi + MTCL2 4LA mut
MTCL2 RNAi + MTCL2 ΔKR mut
%
dendritic length (µm)
D
Maximum of dendritic length (µm)
*
* * *
none none wt ΔKR 4LA
Cont. RNAi MTCL2 RNAi
E
Cont. RNAi
MTCL2 RNAi
none none wt 4LA ΔKR
nuclei
Golgi
dendrite
axon

**Figure S3**. **Effects of MTCL2 knockdown on axonal and dendritic growth of cultured CGNs.**

(A) Percentage of cells with branched axons in the CGNs analyzed in Fig. 2B. The mean values and standard errors of 3 independent experiments are shown for each condition. (B) Distribution of dendritic base numbers among the CGNs analyzed in Fig. 2D. Each data point represents the result of one neuron. Mean values and standard deviations are shown for each scatter plot. (C) Histogram showing the length distribution of all dendrites in the CGNs analyzed in Fig. 2D. Average percentages and standard errors of 3 independent experiments are presented. (D) Maximum dendrite length distribution in the CGNs analyzed in Fig. 2D. Each data point represents the result of one neuron. Mean values and standard deviations are shown for each scatter plot. The paired t-test was used to examine statistical significance (*$p < 0.05$; **$p < 0.01$; n.s., non-significant). (E) Dendritic phenotypes of CGNs subjected to the indicated conditions are illustrated. Note that MTCL2 ΔKR did not rescue any defects in the dendritic extension of MTCL2, whereas MTCL2 4LA showed partial rescue activity against the shortening of preexisting bipolar dendrites, similar to wild-type MTCL2.

### Figure S4

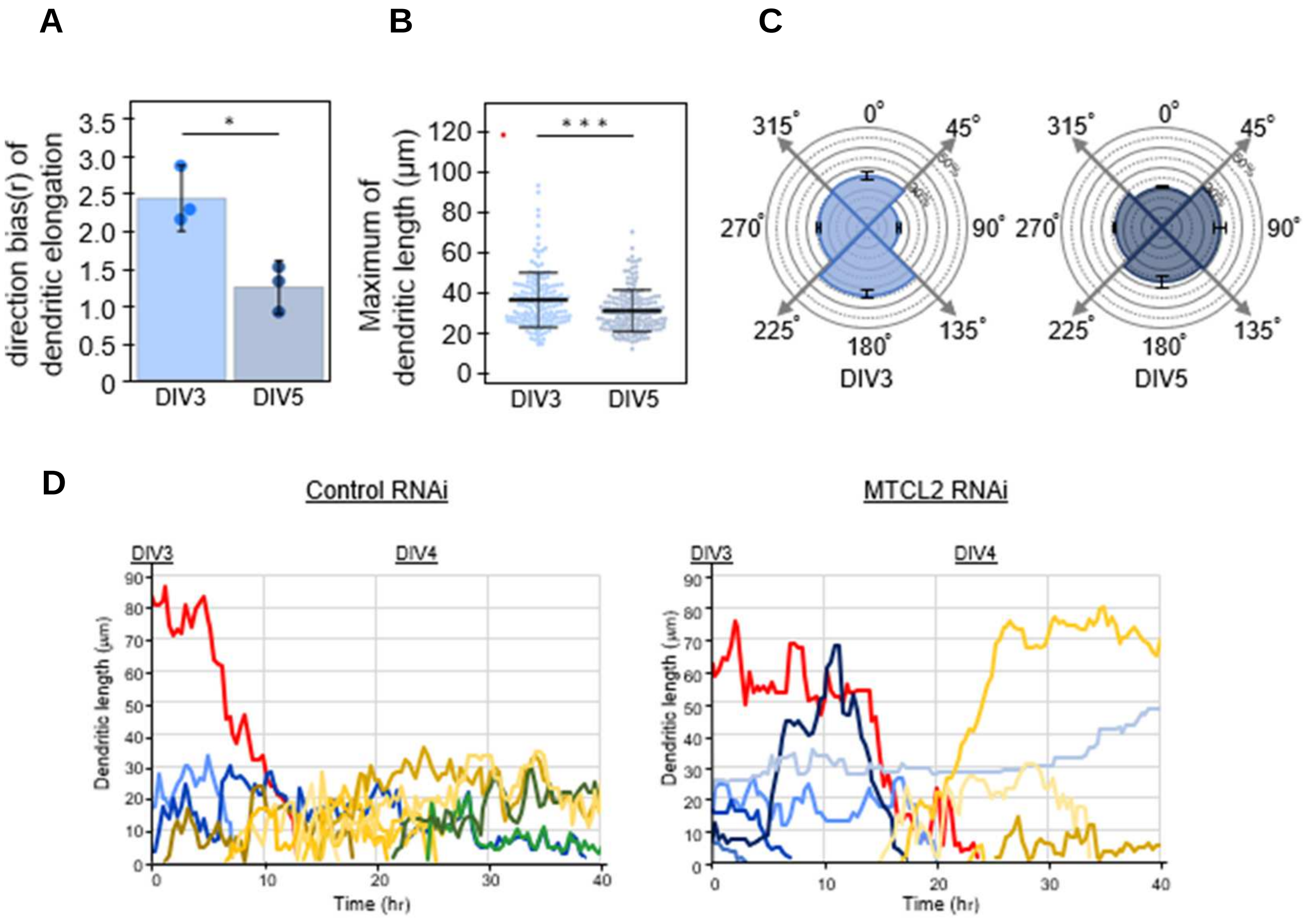


**Figure S4**. **Dendrite extensions of GCNs exhibit bipolar-to-multipolar transitions after DIV3.**

(A) Comparison of the dendritic direction bias (r) between the CGNs at DIV3 and 5 analyzed in Figs. 4B. (B) Maximum dendrite length distributions of the CGNs at DIV3 and 5 analyzed in Figs. 4B. Each data point represents the result of one neuron. Mean values and standard deviations are shown for each scatter plot. (C) θ distribution of the Golgi position in the CGNs at DIV3 and DIV5 analyzed in Fig. 4D. In (A) and (C), the mean values and standard errors of 3 independent experiments are shown. In (A) and (B), the paired t-test was used to examine statistical significance (*$p < 0.05$; ***$p < 0.005$). (D) Dynamic dendrite length changes in the CGNs analyzed in Fig. 4E and F. Data for different dendrites were colored differently. Note that in both cases, the initial long bipolar dendrites exhibited shrinkage. However, the MTCL2-depeleted CGN did not consistently form short dendrites with dynamic features. Instead, the cell re-elongates long dendrites again.

### Figure S5

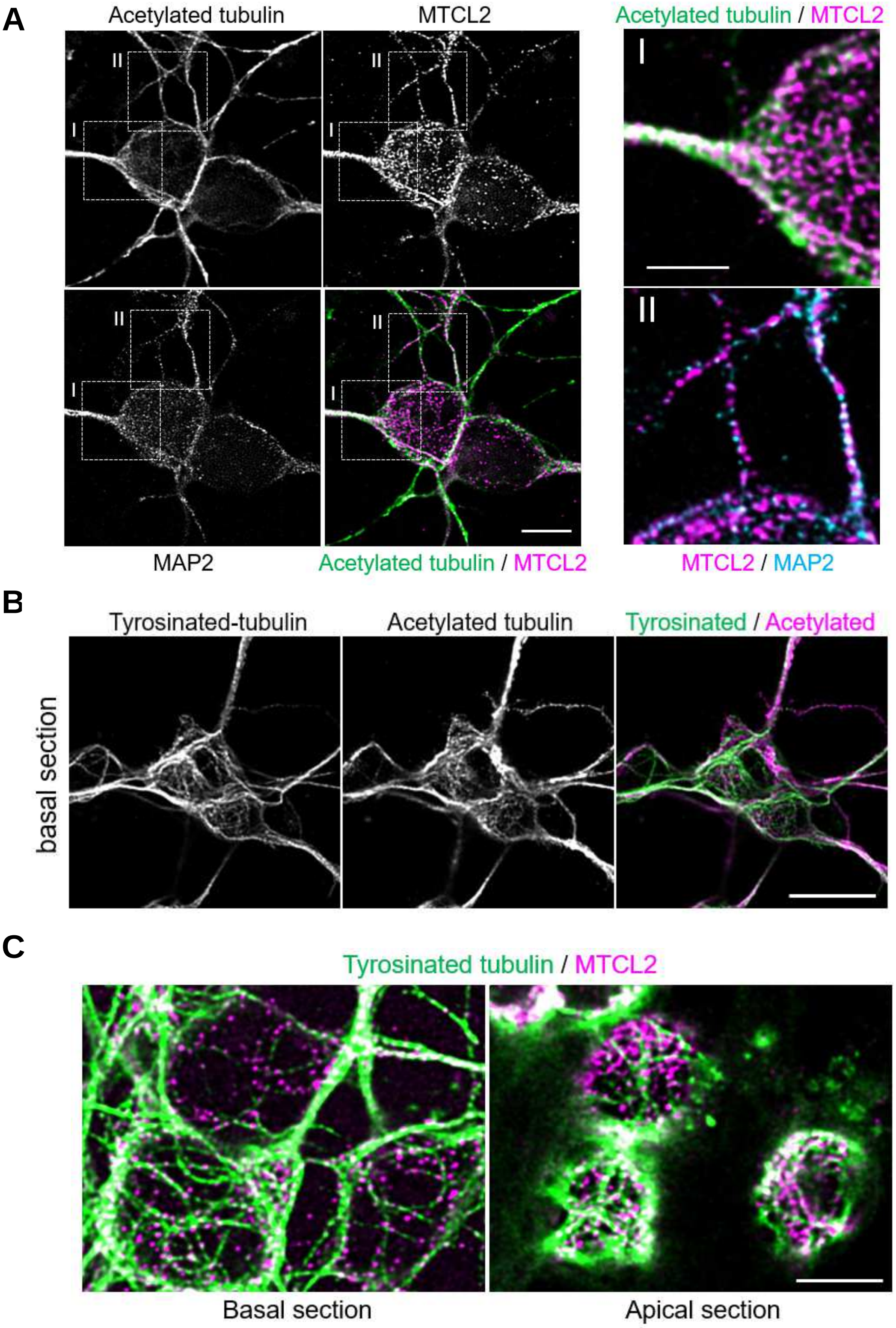
A
Acetylated tubulin
MTCL2
Acetylated tubulin / MTCL2
II
I
MAP2
Acetylated tubulin / MTCL2
MTCL2 / MAP2
B
Tyrosinated-tubulin
Acetylated tubulin
Tyrosinated / Acetylated
basal section
C
Tyrosinated tubulin / MTCL2
Basal section
Apical section

**Figure S5. Colocalization of MTCL2 with MTs in polarized CGNs.**

(A) High-resolution confocal images of CGNs fixed at DIV5 and stained for acetylated tubulin, MTCL2, and MAP2. Single confocal images of the basal plane are shown. The regions indicated by dotted squares (I) and (II) in the left panels are enlarged in the right. Note that, in contrast to MAP2, MTCL2 granules were detected not only on MTs but also in cell bodies. Scale bars: 5 and 2.5 μm in the left and right panels, respectively. (B) High-resolution confocal images of CGNs fixed at DIV5 and stained for tyrosinated and acetylated tubulin. Single confocal images of the basal plane are shown. Note that tyrosinated tubulin is more enriched in the MT network within the cell body than acetylated tubulin. Scale bar: 10 μm. (C) Enlarged merged images of basal (0 μm ) and apical (2.6μm) sections of the CGNs shown in Fig. 5A. Scale bar: 5 μm

### Figure S6

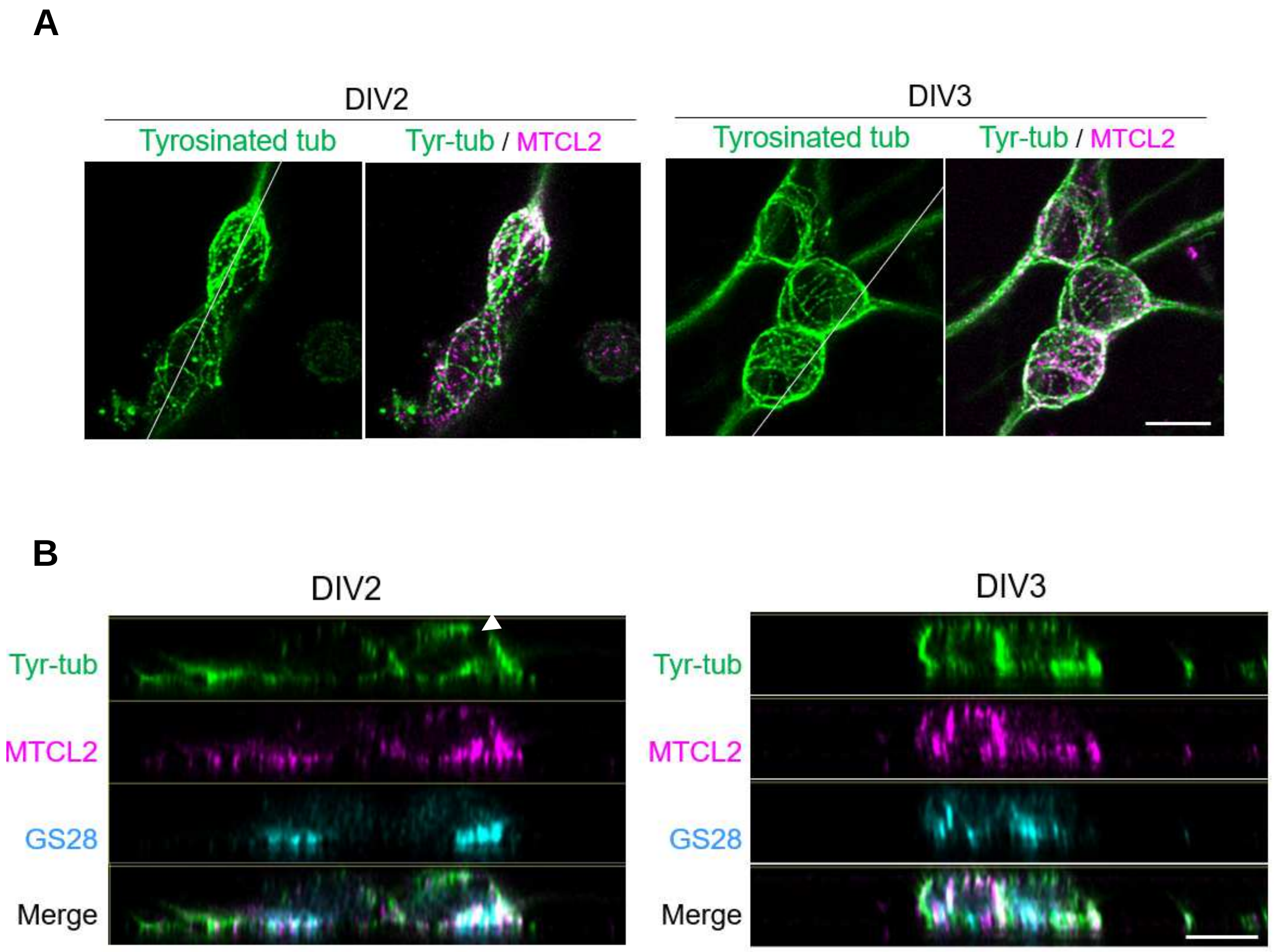


**Figure S6. Colocalization of MTCL2 with MTs in immature CGNs.**

(A) The projected views of the z-stack images of the CGNs analyzed in Fig. 5D. Note that at DIV2, the apical MTs tend to align with the bipolar axis without being organized into the MT cage. At DIV3, an MT cage was detected on which the MTCL2 granules were located. Scale bar: 5 μm. (B) Cross-sectional views of the z-stack images shown in Fig. 5D. The white lines in (A) indicate the position at which each view is created. Scale bar: 5 μm.

### Figure S7

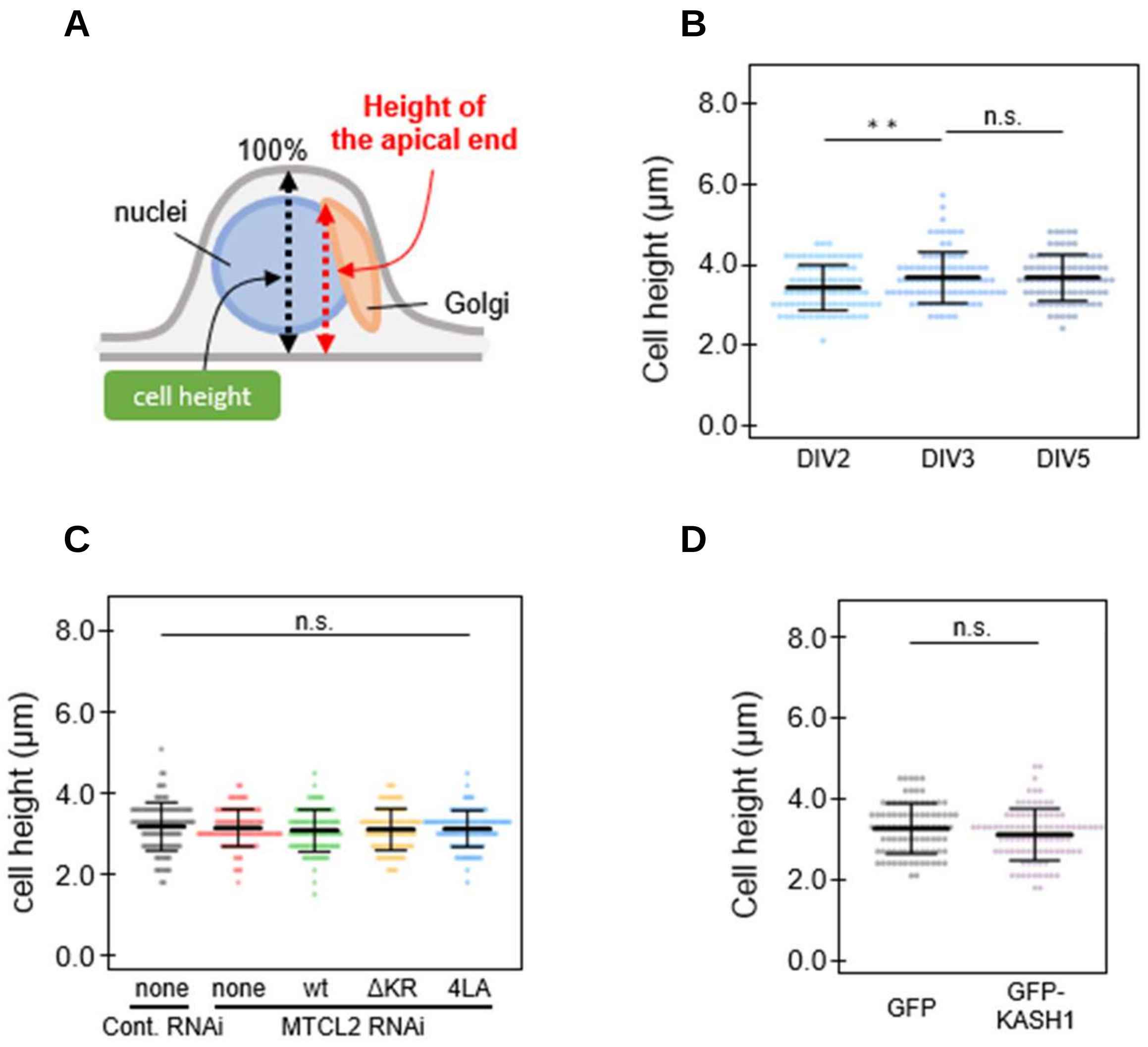


**Figure S7. CGN cell heights at various maturation stages and experimental conditions**

(A) Schematic representation of definition of the cell height and height of the apical edge of the Golgi apparatus measured in this study. (B-D) Height distribution of CGNs at the indicated DIV (B), subjected to the indicated RNAi/rescue conditions (C), or expressing the indicated GFP proteins (D). Each data point represents the result of one neuron observed in 3 independent experiments. Mean values and standard deviations are shown for each scatter plot. The paired t-test was used to examine statistical significance (**$p < 0.01$; n.s., non-significant).

### Figure S8

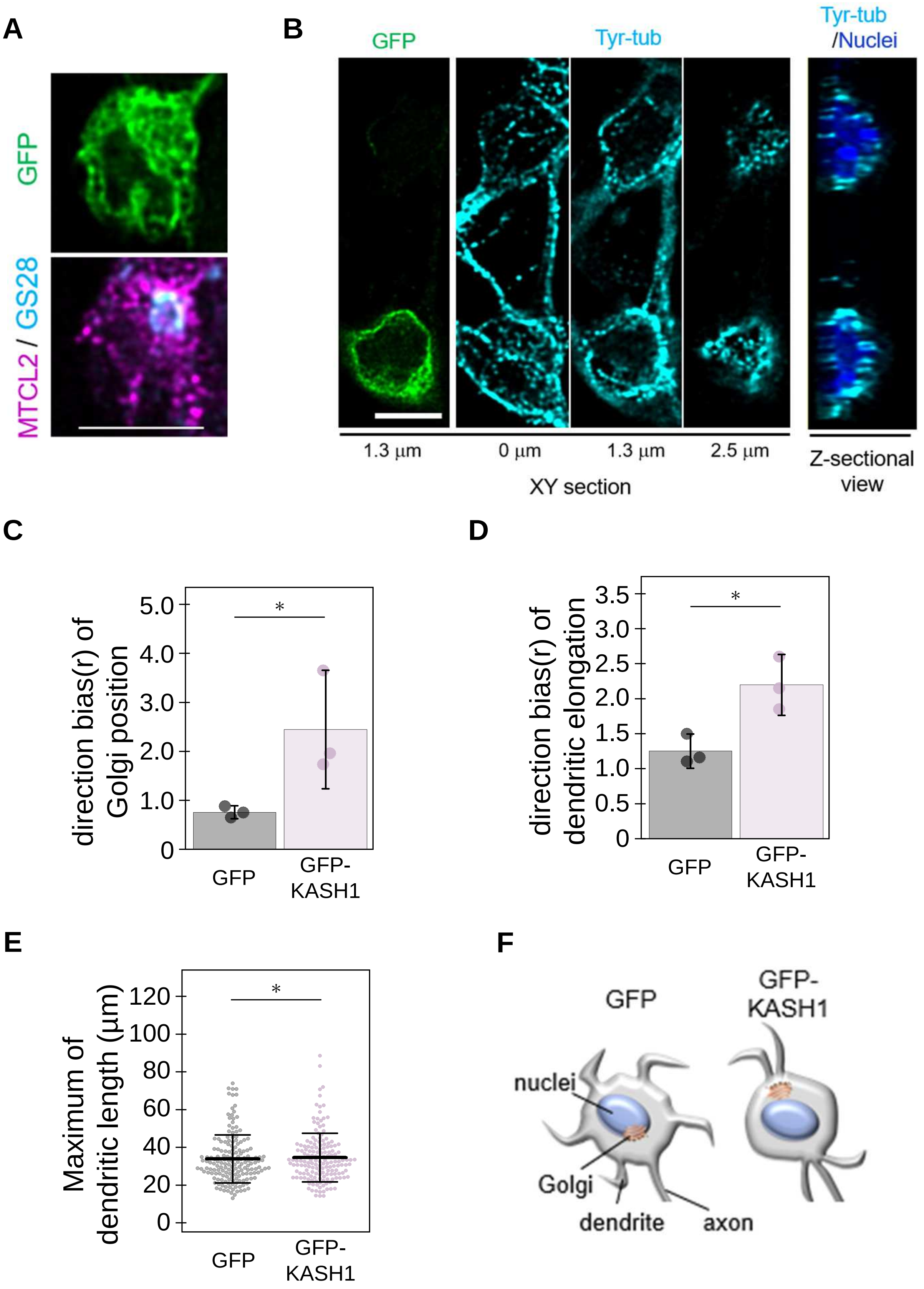

A
GFP
MTCL2 / GS28
B
GFP
Tyr-tub
Tyr-tub
/Nuclei
1.3 μm
0 μm
1.3 μm
2.5 μm
XY section
Z-sectional
view
C
direction bias(r) of
Golgi position
5.0
4.0
3.0
2.0
1.0
0
*
GFP
GFP-
KASH1
D
direction bias(r) of
dendritic elongation
3.5
3.0
2.5
2.0
1.5
1.0
0.5
0
*
GFP
GFP-
KASH1
E
Maximum of
dendritic length (μm)
120
100
80
60
40
20
0
*
GFP
GFP-
KASH1
F
GFP
GFP-
KASH1
nuclei
Golgi
dendrite
axon

**Figure S8. Inhibition of LINC complex activity mimicked most of the phenotypes induced by MTCL2 knockdown.**

(A) A basal confocal section of a GFP-KASH1-expressing cell shown in Fig. 7B. The focus plane is indicated by the dotted lines in Fig. 7B. Scale bar: 5 μm. (B) Serial confocal sections of GFP-KASH1-positive (bottom) and -negative (top) cells from the basal to apical planes are presented. The reconstituted z-sectional images of these cells are displayed on the right. The results of tyrosinated tubulin staining indicated that the formation of the MT-cage was almost unaffected in the GFP-KASH1-expressing cell. Scale bar: 5 μm. (C and D) Comparison of the direction bias (r) of dendrites and the Golgi position between the GFP- and GFP-KASH1-expressing CGNs analyzed in Fig. 7D and F, respectively. The mean values and standard errors of 3 independent experiments are shown for each condition. (E) Distribution of maximum dendrite lengths of the CGNs analyzed in Fig. 7E. Each data point represents the result of a single neuron. Mean values and standard deviations are shown for each scatter plot. For all quantification data, the paired t-test was used to examine statistical significance (*$p < 0.05$). (F) Schematic illustration of the dendritic phenotypes of CGNs expressing GFP and GFP-KASH1.

### Figure S9

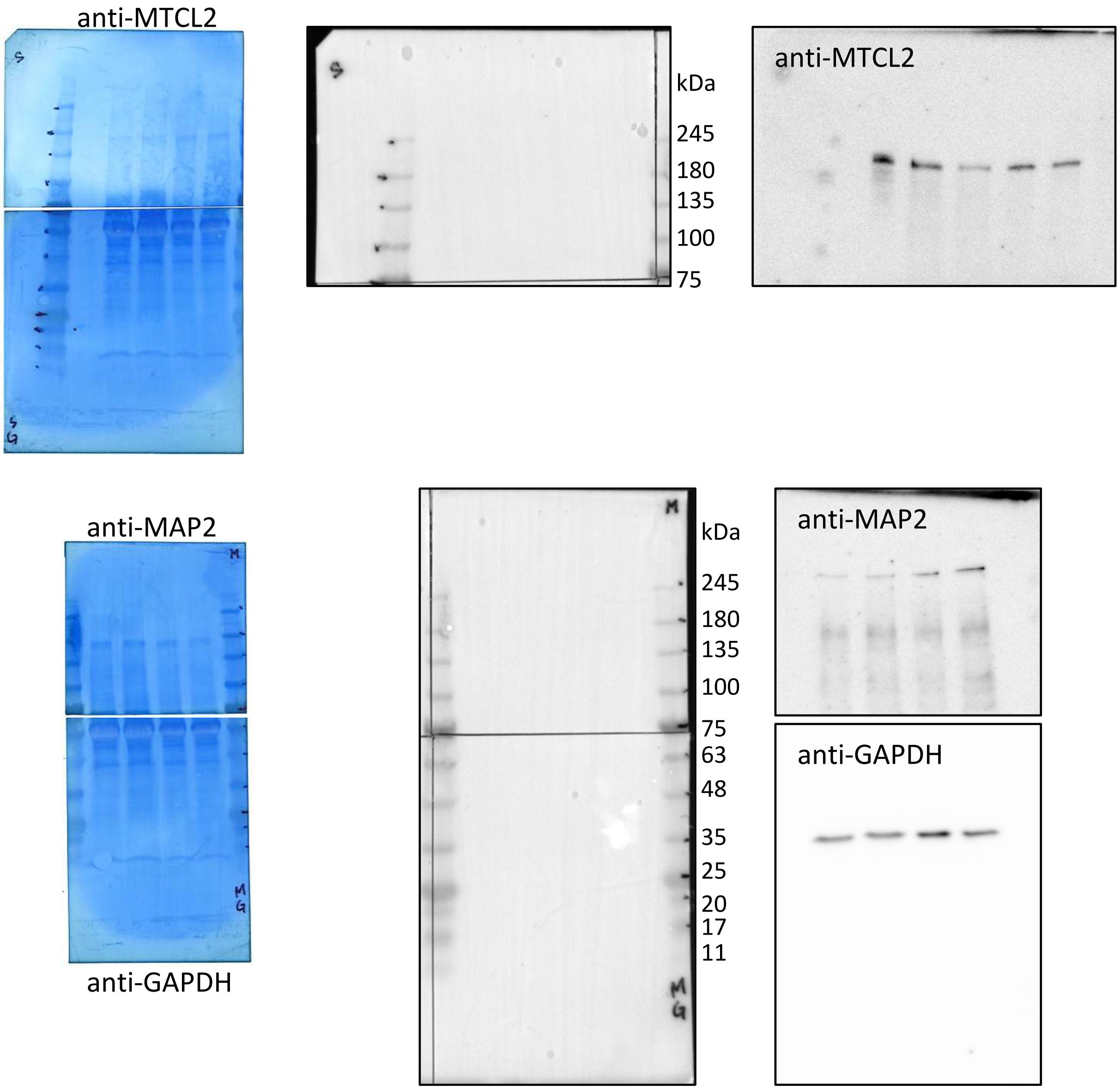


**Figure S9.** Full, uncropped Western blots are shown in Fig. 1F.
